# Identification of lipid senolytics targeting senescent cells through ferroptosis induction

**DOI:** 10.1101/2024.10.14.618023

**Authors:** Lei Justan Zhang, Rahagir Salekeen, Carolina Soto-Palma, Osama Elsallabi, Hongping Ye, Brian Hughes, Borui Zhang, Allancer Nunes, Kyooa Lee, Wandi Xu, Abdalla Mohamed, Ellie Piepgras, Sara J. McGowan, Luise Angelini, Ryan O’Kelly, Xianlin Han, Laura J. Niedernhofer, Paul D. Robbins

## Abstract

Cellular senescence is a key driver of the aging process and contributes to tissue dysfunction and age-related pathologies. Senolytics have emerged as a promising therapeutic intervention to extend healthspan and treat age-related diseases. Through a senescent cell-based phenotypic drug screen, we identified a class of conjugated polyunsaturated fatty acids, specifically α-eleostearic acid and its methyl ester derivative, as novel senolytics that effectively killed a broad range of senescent cells, reduced tissue senescence, and extended healthspan in mice. Importantly, these novel lipids induced senolysis through ferroptosis, rather than apoptosis or necrosis, by exploiting elevated iron, cytosolic PUFAs and ROS levels in senescent cells. Mechanistic studies and computational analyses further revealed their key targets in the ferroptosis pathway, ACSL4, LPCAT3, and ALOX15, important for lipid-induced senolysis. This new class of ferroptosis-inducing lipid senolytics provides a novel approach to slow aging and treat age-related disease, targeting senescent cells that are primed for ferroptosis.

## INTRODUCTION

Aging is the progressive loss of tissue homeostasis, decline in physiological functions and overall health, resulting in an increased prevalence of numerous chronic diseases. This decline is driven by several cellular and molecular hallmarks, including genomic instability, telomere attrition, epigenetic alterations, loss of proteostasis, deregulated nutrient sensing, mitochondrial dysfunction, cellular senescence, stem cell exhaustion, and altered intercellular communication.^1–3^ These hallmarks of aging are not isolated events, but instead are interconnected, with changes in one often influencing many others. Among them, cellular senescence stands out as a critical hallmark, as genetic and pharmacologic approaches have intricately linked the aged-dependent increase in the senescent cell burden to nearly all other pillars of aging.^4,5^ Cellular senescence is characterized by a stable cell cycle arrest and upregulation of senescent cell anti-apoptotic pathways (SCAPs) despite remaining metabolically active.^6,7^ Another defining feature of senescent cells is the development of a senescence-associated secretory phenotype (SASP), which includes bioactive molecules such as chemokines, cytokines, proteases, growth factors, metabolites, nucleic acid, and extracellular vesicles.^8^ Senescence can be triggered by various cellular stressors including genotoxic damage, oxidative stress, replicative exhaustion, oncogenic signals, and viral infections.^9^ This intricate interplay with the hallmarks of aging conferred by the SASP positions cellular senescence as a key driver of aging.

Senescent cells persist in tissues due to their exit from the cell cycle and resistance to cell death. As these cells accumulate with age, they can reach a critical threshold where they detrimentally impact the surrounding microenvironment through their sustained expression SASP, thereby contributing to the pathophysiology of aging and related disorders.^10–12^ The role of senescent cells in driving various age-related diseases, including diabetes, cancer, osteoarthritis, and Alzheimer’s disease, has been established in transgenic mouse models where certain types of senescent cells (e.g., p16^INK4a+^ and p21^Cip1+^) can be ablated.^13–17^ More importantly, pharmacological targeting senescent cells with senolytics, agents that selectively eliminate these cells, has emerged as a promising strategy for extending healthspan and treating many age-related diseases.^18–20^ However, many of the identified senolytics are anti-cancer agents that have associated adverse effects. As such, there is growing interest in developing more effective and safer senolytics.^9,21,22^

Fatty acids are naturally occurring compounds found in various food sources, including plants, animals, and microorganisms. Many fatty acids have been reported to have diverse therapeutic effects on human health.^23^ However, whether the benefits of certain fatty acids are conferred, at least in part, through senotherapeutic activity, remains unexplored. In this study, we employed a senescent cell-based phenotypic drug discovery approach to screen a focused library of diverse fatty acids and identified a novel class of polyunsaturated fatty acids with senolytic properties both in cell culture and in multiple mouse models of aging. Mechanistically, we show that these effects are conferred through selective ferroptosis induction in senescent cells which are primed with upregulated cellular iron, ROS levels, cytosolic PUFAs, and key enzymes promoting iron-dependent cell death.

## RESULTS

### Phenotypic screening identifies potent lipid senolytics

To identify compounds with senotherapeutic activity, we developed previously a robust phenotypic drug screening platform based on the elevated senescence-associated beta-galactosidase (SA-β-gal) activity in senescent cells.^9,11^ This platform enables the screening of compounds across various cell types where senescence can be induced using different methods. Briefly, the senescent cells are seeded into 96-well plates and uniformly treated with test compounds with non-senescent cells used as a control. Post-treatment, the percent of SA-β-Gal-positive senescent cells is assessed using the fluorogenic dye C_12_FDG.^24^ This lipophilic dye produces a strong fluorescent signal upon cleavage by β-galactosidase and is a used to detect SA-β-gal activity in senescent cells at pH 6.^25^ This platform is equipped with a high-content fluorescent imaging system and allows rapid evaluation of two categories of senotherapeutics: senolytics, which selectively kill senescent cells and reduce the number of C_12_FDG-positive cells, and senomorphics, which reduce the senescence phenotype without inducing cell death (Figure 1A).

**Figure 1.**
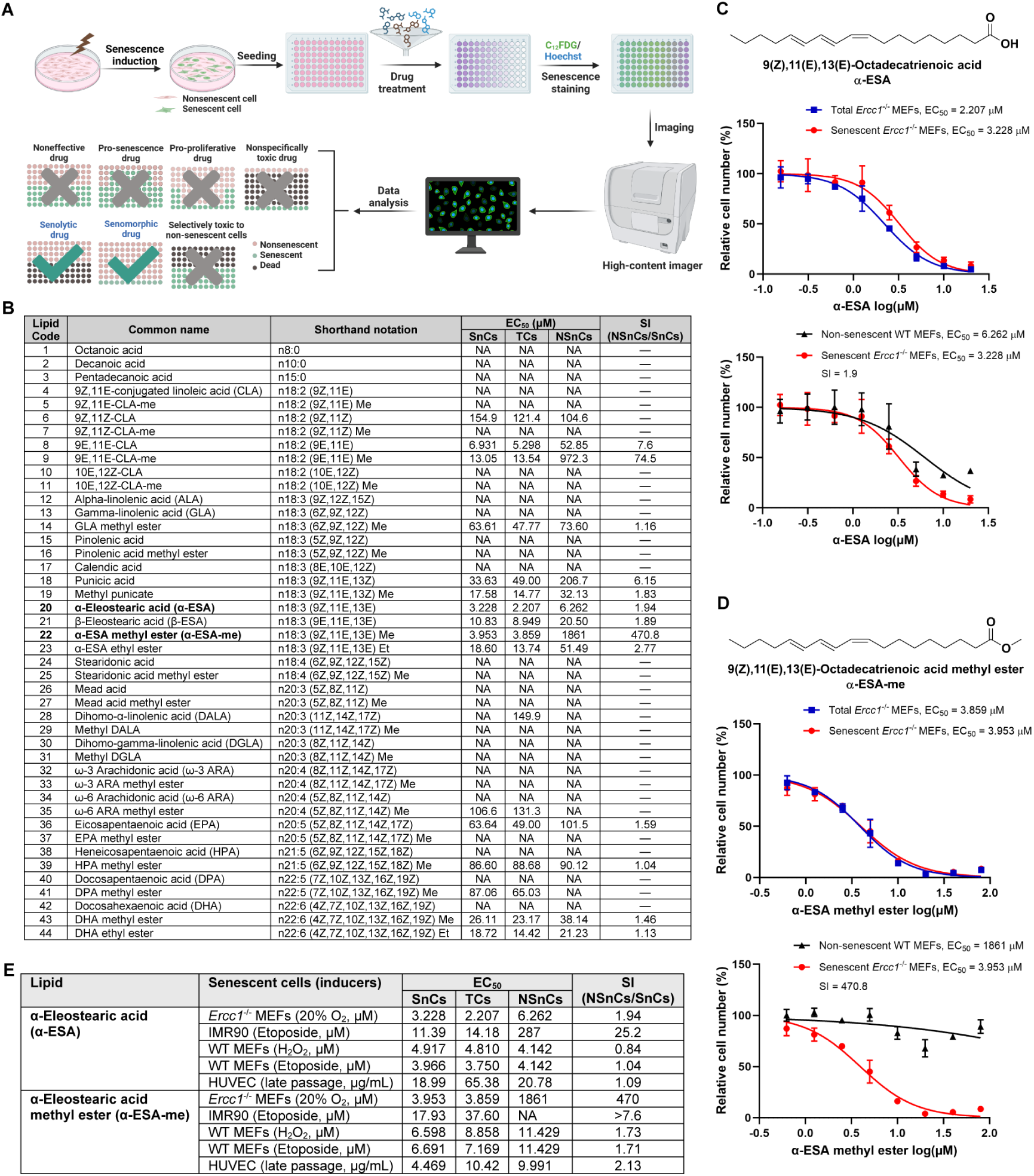
Identification of novel lipid senolytics through senescent cell-based phenotypic drug screening. (A) Schematic diagram of the senescent cell-based phenotypic drug screening platform for identifying novel senotherapeutics. *Created in BioRender.com* (B) Summary of senescent cell screening of various fatty acids in non-senescent WT MEF and senescent *Ercc1*^-/-^ MEF cells. Abbreviations: **SnCs**: C_12_FDG-positive senescent cells; **TCs**: Total *Ercc1*^-/-^ MEF cells; **NSnCs**: Non-senescent WT MEF cells; **SI**: Selectivity index. **EC_50_**: Concentration resulting in a 50% reduction in cell number compared to control. **NA**: EC_50_ not available due to inactivity. (C) Chemical structure of α-ESA and its dose-response curves for senolytic EC_50_ determination. Error bars represent SD for n = 3. (D) Chemical structure of α-ESA-me and its dose-response curves for senolytic EC_50_ determination. Error bars represent SD for n = 3. (E) Overview of the senolytic effects of α-ESAs across a broad range of senescent cell types.

Using this screening platform, we conducted a focused screen of a panel of various fatty acids reported to have health benefits and dietary functions.^23^ The panel included medium-chain fatty acids with aliphatic tails of 6 to 12 carbons, long-chain fatty acids with tails of 13 to 21 carbons, and very long-chain fatty acids with tails of 22 or more carbons (Figure 1B). We particularly focused on conjugated fatty acids (CFAs), such as polyunsaturated fatty acids (PUFAs) like eicosapentaenoic acid (EPA) and docosahexaenoic acid (DHA),^26,27^ which tend to exhibit stronger physiological effects compared to non-conjugated fatty acids.^28^

Initially, we used primary *Ercc1*^-/-^ mouse embryonic fibroblasts (MEFs) for our screen, which were induced into senescence through oxidative stress by passaging at 20% O_2_.^29^ The ERCC1 protein, a crucial component of the ERCC1/XPF endonuclease complex, plays an essential role in repairing multiple types of DNA damage. Mice with reduced ERCC1 expression (*Ercc1*^-/Δ^) exhibit increased sensitivity to DNA damage, an increased accumulation of senescent cells, accelerated aging phenotypes, and a reduced lifespan.^30^ Accordingly, *Ercc1*^-/-^ MEFs are impaired in DNA damage repair and thus are prone to undergoing senescence when subjected to oxidative stress when cultured at elevated oxygen levels. Wild-type (WT) primary MEF cells were used as the non-senescent, proliferative control. Both cell types were treated with the lipid panel for 48 hours, followed by staining with C_12_FDG to detect SA-β-gal+ cells. Each lipid was tested across a range of concentrations to determine the EC_50_ values, the effective concentration needed to achieve a 50% reduction in cell number compared to the control (Figure 1B). The senolytic and senomorphic effects were evaluated by comparing the EC_50_ values of C_12_FDG-positive cells and the total number of senescent cells. The selectivity index (SI) was calculated by the ratio of EC_50_ values between non-senescent cells (NSnCs) and senescent cells (SnCs).

Among the diverse lipids tested, α-eleostearic acid (α-ESA, lipid 20, EC_50_ = 3.228 µM, SI = 1.94) and α-ESA methyl ester (α-ESA-me, lipid 22, EC_50_ = 3.953 µM, SI = 470) exhibited the strongest senolytic activity (Figures 1B-D). Other lipids, such as 9E, 11E-conjugated linoleic acid (CLA, lipid 4) and its methyl ester (lipid 5) had slightly weaker senolytic activity, while punicic acid (lipid 18), methyl pumicate (lipid 19), DHA esters (lipids 43, 44), β-ESA (lipid 21) and α-ESA ethyl ester (lipid 23) showed no to weak senolytic activity at the concentration ranges tested.

### Structure-activity relationship (SAR) analysis reveals key structural features for senolytic activity

In general, medium-chain fatty acids and saturated fatty acids demonstrated no senolytic potential, as exemplified by octanoic acid, decanoic acid, and pentadecanoic acid (lipids 1-3). Although certain long-chain fatty acids in the n20 and n22 series, including arachidonic acid (lipid 32), EPA (lipid 36), HPA (lipid 38), DPA (lipid 40), and DHA (lipid 42), are known for their important dietary functions,^26,27^ they exhibited little to no senolytic activity. This suggests that simply increasing chain length alone does not necessarily enhance senolytic activity. Notably, several long-chain fatty acids with 18 carbon tails demonstrated stronger senolytic activities than those with other chain lengths. For example, 9Z,11E-conjugated linoleic acid (CLA, lipid 4) with two conjugated double bonds, punicic acid (lipid 18), and α-ESA (lipid 20) with three conjugated double bonds exhibited stronger senolytic activity.

In addition to carbon chain length, the position and configuration of double bonds in fatty acids are also critical for the senolytic activity. For instance, CLAs can exist in either *cis* or *trans* configurations, and our screening found that the 9E,11E-CLA isomer (lipid 8) is a highly potent and selective senolytic, whereas the 9Z,11E-CLA (lipid 4), 9Z,11Z-CLA (lipid 6), and 10E,12Z-CLA (lipid 10) isomers are inactive. The configuration effect was also observed in the ESA series, where α-ESA (lipid 20) with a 9Z,11E,13E configuration is more potent than its all-*trans* counterpart β-ESA (9E,11E,13E, lipid 21).

Interestingly, esterification appears to be a key modification that can alter the senolytic activity and selectivity of fatty acids. For example, while unmodified DHA (lipid 42) showed no significant activity, its methyl and ethyl esters (lipids 43 and 44) displayed some senolytic properties. This trend also was observed with several other fatty acids, where esterification improves selectivity, albeit sometimes at the expense of potency. For instance, 9E,11E-CLA-Me (lipid 9) is less potent but 10-fold more selective than 9E,11E-CLA (lipid 8). An exception is α-ESA methyl ester (α-ESA-me, lipid 22), which exhibited a striking selectivity index of 470, though less potent, compared to its non-esterified form α-ESA (lipid 20), which has a much lower selectivity index of 1.94 (Figures 1C and 1D).

Another major distinction between the active and inactive unsaturated fatty acids is the presence of conjugation. Unconjugated fatty acids, even those with multiple double bonds, generally did not exhibit senolytic properties. For example, in the n18 series of octadecanoids, all the inactive unsaturated fatty acids are methylene-interrupted polyenes, such as alpha-linolenic acid (ALA, lipid 12), gamma-linolenic acid (GLA, lipid 13), pinolenic acid (lipid 15), and stearidonic acid (lipid 24). This pattern extends to the n20 series (e.g., mead acid or lipid 26, DALA or lipid 28, DGLA or lipid 30, arachidonic acids or lipids 32 and 34, EPA or lipid 36), the n21 series (e.g., HPA or lipid 38), and the n22 very long-chain fatty acids (e.g., DPA or lipid 40, DHA or lipid 42), regardless of whether they are omega-3 or omega-6 fatty acids.

Collectively, these SAR findings underscore the importance of molecular structure in determining the senolytic potency and selectivity of fatty acids. From this initial screening, we selected the most selective senolytic, α-ESA-me (lipid 22, EC_50_ = 3.953 µM, SI = 470), and the most potent senolytic, α-ESA (lipid 20, EC_50_ = 3.228 µM, SI = 1.94), for further investigation.

### α-ESAs are novel lipid senolytics that eliminate diverse set of senescent cells

To further validate the senolytic activity of α-ESA and its derivative α-ESA-me, we used several additional senescent cell models. These included WT MEF cells induced to senescence by oxidative stress (H_2_O_2_) and genotoxic stress (etoposide), human IMR90 fibroblast cells induced by genotoxic stress (etoposide), and human umbilical vein endothelial cells (HUVECs) induced by replicative passaging. Results showed that both α-ESA and α-ESA-me exhibited significant senolytic activity, able to effectively reduce C_12_FDG-positive senescent cells across a broad range of senescent cell types and inducers (Figure 1E and S1A). While α-ESA demonstrated greater potency, α-ESA-me exhibited higher selectivity, particularly in *Ercc1*^-/-^ MEFs. Interestingly, both compounds showed comparable selectivity in IMR90 cells, (Figure S1B), indicating some cell type-specific variations in their senolytic activities.

To further explore the senolytic actions of α-ESA and α-ESA-me, we examined their effects over time ranging from 12 to 48 hours (Figure S1C). The EC_50_ of the senolytic effect of α-ESA improved from 5.568 µM at 12 hours to 1.330 µM at 36 hours. However, its senolytic EC_50_ value rose slightly to 2.195 µM by 48 hours. In contrast, the EC_50_ value of α-ESA-me decreased over time, showing less potency than α-ESA, but offering a more sustained effect across all time points. These initial results suggest that α-ESA acts more rapidly to induce senolysis whereas α-ESA-me is more stable, providing a longer-lasting senolytic effect.

### α-ESAs reduce tissue senescence and extend healthspan in aged mice

To evaluate the senolytic potential of the lipid senolytics α-ESA and α-ESA-me *in vivo*, we tested these compounds in 20-22 months old WT C57BL/6 mice. The mice were treated with 50 mg/kg of either α-ESA or α-ESA-me for five consecutive days. Two days after the last administration, the mice were sacrificed for tissue analysis (Figure 2A). Molecular analysis revealed that α-ESA-me had a superior effect in reducing tissue senescence compared to α-ESA at this dosage, particularly in the liver and heart, while α-ESA also demonstrated efficacy in reducing senescence in the brain (Figures 2B and S2A). Further studies on 32-month-old C57BL/6 mice treated under the same regimen (Figure 2C) showed that α-ESA-me significantly reduced senescence and SASP factors in multiple tissues, including genes encoding p16^INK4a^, p21^Cip1^, TNFα, IL-6, CXCL1, MCP1, and IL-1β (Figures 2D and S2B). The reduction was particularly pronounced in the kidney (Figure 2E), liver, and lung (Figure S2B). Additionally, α-ESA-me treatment led to a marked decrease in the proportion of p21^Cip1^-positive gamma delta (γδ) T cells in the spleen, suggesting a possible senolytic effect on specific senescent immune cells in aged mice (Figure 2F).

**Figure 2.**
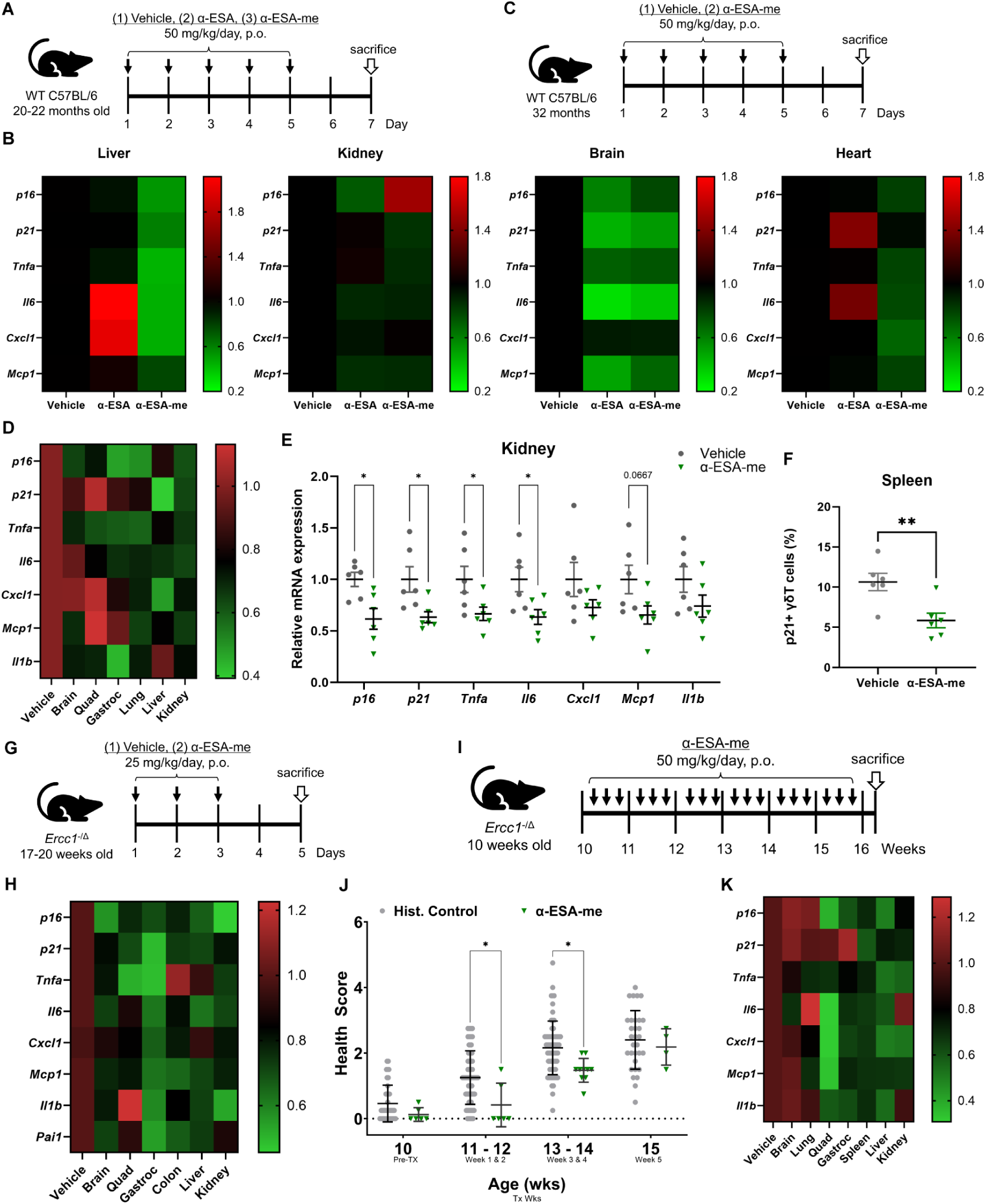
Evaluation of the senolytic effects of α-ESA and α-ESA-me in models of accelerated aging and naturally aged mice. (A) WT C57BL/6 mice (20-22 months old) were treated with 50 mg/kg of α-ESA or α-ESA-me by oral gavage for five consecutive days. Tissues were collected two days after the last dose for analysis. (B) Effects of α-ESA and α-ESA-me on reducing senescence across different tissues in aged WT C57BL/6 mice. (C) WT C57BL/6 mice (32 months old) were treated with 50 mg/kg of α-ESA-me by oral gavage for five consecutive days. Tissues were collected two days after the last dose for analysis. (D) Reduction of senescence markers by α-ESA-me in multiple tissues of 32-month-old WT mice, particularly in (E) the kidney. Error bars represent SEM for n = 6. (F) α-ESA-me reduced p21+ senescent γδ T cells in the spleen of the 32-month-old WT C57BL mice. Error bars represent SEM for n = 6. (G) *Ercc1*^-/Δ^ progeria mice (17-20 weeks old) were treated with 25 mg/kg of α-ESA-me for three consecutive weeks by oral gavage. Mice were sacrificed two days after the last dose. (H) Acute α-ESA-me treatment reduced senescence in multiple tissues of *Ercc1*^-/Δ^ progeria mice. (I) *Ercc1*^-/Δ^ progeria mice (10 weeks old) were treated with 50 mg/kg of α-ESA-me for six consecutive weeks by oral gavage. Weekly health assessments were conducted to score age-related symptoms, including tremor, kyphosis, dystonia, ataxia, gait disorder, hindlimb paralysis, and forelimb grip strength. (J) Chronic α-ESA-me treatment significantly improved composite health scores in *Ercc1*^-/Δ^ progeria mice. (K) Chronic α-ESA-me treatment reduced tissue senescence and SASP factors in *Ercc1*^-/Δ^ progeria mice. Error bars represent SEM for n = 6 (vehicle) and n = 4 (α-ESA-me).

Given the stronger senolytic effect of α-ESA-me compared to α-ESA in naturally aged mice, we selected α-ESA-me for further evaluation on healthspan in *Ercc1*^-/Δ^ progeria mice, a well-established model of accelerated senescence and aging.^31^ Initially, to validate the senolytic effect of α-ESA-me in this model, we performed a short-term, acute treatment of *Ercc1*^-/Δ^ mice with α-ESA-me for three days (Figure 2G). RT-qPCR analysis of different tissues revealed a notable decrease in senescence markers and SASP factors, particularly in the kidney, liver, and gastrocnemius muscles (Figures 2H and S3A).

To assess the long-term effects of α-ESA-me on healthspan, we treated the *Ercc1*^-/Δ^ mice with α-ESA-me orally three times per week for six weeks, starting at 10 weeks of age (Figure 2I). Weekly health assessments were conducted to monitor age-related symptoms, including tremor, kyphosis, dystonia, ataxia, gait disorder, hindlimb paralysis, and forelimb grip strength. The results indicated that α-ESA-me effectively reduced the composite score of aging symptoms without negatively affecting body weight, consistent with a lack of toxicity (Figures 2J and S3B). In particular, it improved tremor and kyphosis in treated mice (Figure S3B). Although no significant reduction in overall aging symptoms was observed by week 16 (Figure 2J), RT-qPCR analysis of tissues continued to show a noticeable decrease in senescence and SASP markers in kidney, liver, spleen, and muscle (Figures 2K and S3C). These results further confirm the senolytic activity of α-ESA-me.

### α-ESAs induce senescent cells death *via* ferroptosis

To elucidate the mechanism through which α-ESA and α-ESA-me induce senescent cell death, we first investigated whether the metabolites of α-ESAs contribute to the senolytic activity. α-ESA has been reported to be rapidly converted to conjugated linoleic acid^32^ and further metabolized into gamma linolenic acid and arachidonic acid through Δ6 and Δ5 desaturase, respectively (Figure S4E). To test whether these metabolic pathways were involved in the senolytic effects of α-ESAs, we pretreated senescent cells with inhibitors of Δ6 desaturase (SC-26196) and Δ5 desaturase (sesamin) prior to the lipid senolytic treatment. The results showed that neither SC-26196 nor sesamin prevented the senolytic effects of α-ESA-me (Figures 3A) or α-ESA (Figure S4C). Similar results were observed in senescent cells induced by either oxidative or genotoxic stress (Figures S4A and S4B). These results suggest that the observed senescent cell death is not mediated through metabolic conversion of the α-ESAs.

**Figure 3.**
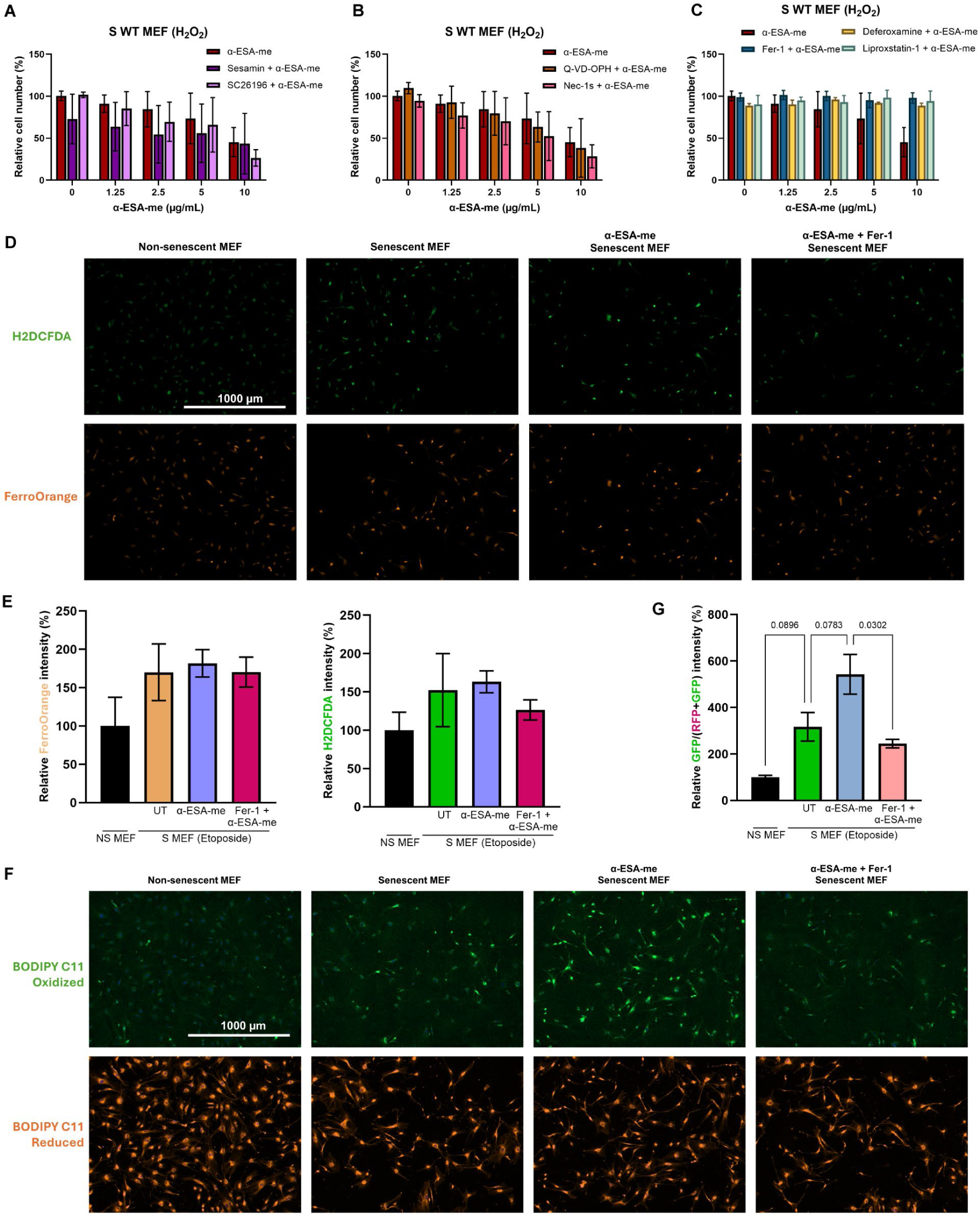
α-ESAs induced senolysis is mediated by ferroptosis. (A-C) Senescent MEF cells were treated with α-ESA-me for 48 hours, with or without 1- hour pretreatment with the following compounds: sesamin (50 μM), SC26196 (200 nM), Q-VD-OPH (20 μM), Nec-1s (50 μM), Fer-1 (2 μM), deferoxamine (50 μM), or liproxstatin- 1 (2 μM). Error bars represent SD for n = 3. (D) Cells were treated with α-ESA-me (5 µg/mL) for 6 hours, with or without 1-hour pretreatment with Fer-1 (2 μM). Ferrous iron and ROS levels were detected by FerroOrange and H2DCFDA, respectively. (E) Quantification of the ferrous iron and ROS levels. Error bars represent SD for n = 2. (F) Lipid peroxidation was detected by C11 BODIPY after treatment with α-ESA-me (5 µg/mL) for 6 hours, with or without Fer-1 (2 μM). (G) Quantification of the lipid peroxidation. Error bars represent SD for n = 2.

Since most existing senolytics function by inducing apoptosis,^21^ we next examined whether α-ESA and α-ESA-me triggered senescent cell death *via* apoptosis. Surprisingly, pretreatment with Q-VD-OPH, an irreversible pan-caspase inhibitor, did not protect senescent MEFs from cell death induced by either α-ESA or α-ESA-me (Figures 3B and S4A-C). Similarly, the necroptosis inhibitor Nec-1s also failed to prevent senescent cell death. However, pretreatment with the ferroptosis inhibitor Fer-1 completely blocked the senolytic effects of α-ESA and α-ESA-me (Figures 3C and S4A-C). Further investigation using other ferroptosis pathway regulators, such as the iron chelator deferoxamine and the lipid peroxidation inhibitor liproxstain-1, confirmed that inhibiting ferroptosis-related pathways prevented senescent cell death induced by the α-ESAs (Figures 3C and S4A-C). These results suggest that ferroptosis, rather than apoptosis or necroptosis, is the primary mode of cell death induced by these novel lipid senolytics.

Ferroptosis is an iron-dependent form of programed cell death triggered by the accumulation of reactive oxygen species (ROS) and lipid peroxidation.^33^ We found that senescent cells exhibited higher levels of ferrous iron and ROS compared to non-senescent cells, as detected by FerroOrange and H_2_DCFDA, respectively (Figures 3D and 3E). α-ESA-me treatment did not significantly alter iron or ROS levels in the cells. However, the addition of the ferroptosis inhibitor Fer-1 reduced ROS levels, but not ferrous iron (Figure 3E). Lipid peroxidation, another key factor in ferroptosis, was elevated in senescent cells compared to non-senescent cells (Figures 3F and 3G). Notably, α-ESA-me further exacerbated this lipid peroxidation, whereas ferroptosis inhibitor Fer-1 blocked this effect. A similar ferroptosis-dependent effect also was observed with α-ESA treatment (Figures S4D).

These results suggest that ferroptosis is a novel mechanism of senescent cell death. To further validate this, we treated senescent cells with several ferroptosis inducers, such as the GPX4 inhibitor RSL3 and the glutamate-cystine antiporter Xc inhibitor erastin. Indeed, these treatments resulted in moderate senolysis across various senescent cell models, regardless of cell types or senescence inducers (Figure S5). Collectively, these findings strongly suggest that the senescent cell death induced by α-ESA and α-ESA-me occurs through ferroptosis, unlike the conventional apoptotic senescent cell death.

### α-ESAs promote lipid peroxidation in senescent cells in a ferroptosis-dependent manner

To examine the effects of α-ESAs on the lipidome, we conducted an undirected shotgun lipidomic analysis. Non-senescent (NS), H_2_O_2_-induced (S-H_2_O_2_), and etoposide-induced senescent (S-Eto) MEF cells were treated with the α-ESAs for eight hours and then collected for lipidomic analysis before overt cell death occurred. A total of 310 lipid species across 20 classes were identified (Table S1), with free cholesterol (FC), phosphatidylcholine (PC), and phosphatidylethanolamine (PE) being the most abundant lipid species (Figure 4A). Class wise comparisons of mean Z-scores across species revealed distinct lipidomic signatures between NS and S-Eto MEFs (Figure 4B). Untreated senescent MEFs showed substantial downregulation of many lipid classes, including triacylglycerols (TAGs), phosphatidic acid (PA), phosphatidylinositol (PI), and phosphatidylserine (PS), compared to NS MEFs (Figure 4B). The S-Eto MEFs in turn exhibited upregulated levels of bis(monoacylglycero) phosphate (BMP) and phosphatidylglycerol (PG). At the species level, untreated S-Eto MEFs had strong upregulation of lyso-phosphatidylcholine (LPC), BMP and lyso-phosphatidylethanolamine (LPE) with stearic, arachidonate, linoleic, palmitoleic and behenic residues (Figure 4C).

**Figure 4.**
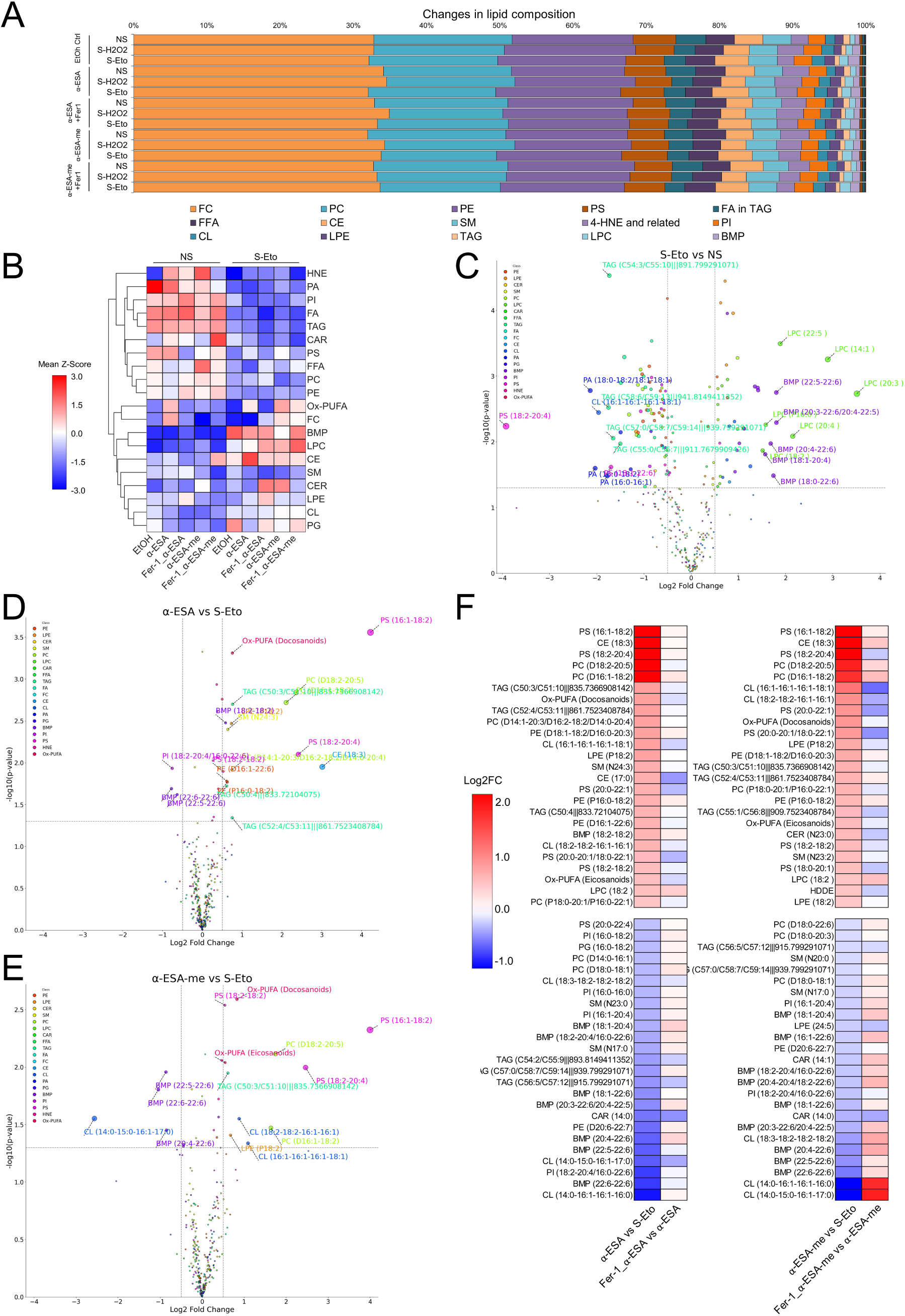
Lipidomic analysis of cells treated with α-ESAs. Non-senescent (NS) and etoposide (S-Eto)/H_2_O_2_-induced (S-H2O2) senescent MEF cells were treated with α-ESA (2 µg/mL) or α-ESA-me (2 µg/mL) for 8 hours, with or without 1- hour pretreatment of Fer-1 (2 μM). (A) Stacked bar graph displaying the distribution of lipid species. The mol% represents the moles of each lipid species extracted from a sample, calculated as a percentage of the total lipid extract. Data are based on n= 4 for all groups. (B) Euclidian clustered heatmap showing color-coded Z-scores for lipid class-wise distribution. Data are based on mean Z-scores for species within each class and n= 4 for all groups. (C) Volcano plot of differential changes in abundance of lipid species for S-Eto vs NS groups; lipid species with log2FC>|1.5| and p<0.05 are labeled; colors represent lipid classes. (D-E) Volcano plot of differential changes in abundance of lipid species for α-ESA vs S-Eto and α-ESA-me vs S-Eto groups; lipid species with log2FC>|0.5| and p<0.05 are labeled; colors represent lipid classes. (F) Heatmap of top 25 upregulated and downregulated lipid species with α-ESA and α-ESA-me treated MEFs compared to S-Eto. Adjacent column shows change in abundance in Fer-1 pretreated vs α-ESA and α-ESA-me treated MEFs. Abbreviations: PE, Phosphatidylethanolamine; LPE, Lyso-Phosphatidylethanolamine; CER, Ceramide; CE, Cholesterol Esters; SM, Sphingomyelin; PC, Phosphatidylcholine; LPC, Lyso-Phosphatidylcholine; CAR, Acyl-Carnitine; FFA, Free Fatty Acid; TAG, Triacylglycerol; FA, Fatty Acyl Chains in TAG; CL, Cardiolipin; PA, Phosphatidic acid; PG, Phosphatidylglycerol; BMP, Bis(Monoacylglycero) Phosphate; PI, Phosphatidylinositol; PS, Phosphatidylserine; Ox-PUFA, Oxidized Polyunsaturated Fatty Acids.

These changes in abundance could reflect membrane phospholipid disintegration and cleavage in the senescent cells. These also suggest that senescent cells, in addition to ferrous iron and ROS, have a higher abundance of lipid peroxidation substrates, such as cytosolic PUFAs, priming the cells for ferroptosis initiation. Senescent cells did not exhibit substantially elevated levels of 4-hydroxy-2-nonenal (HNE) or oxidized polyunsaturated fatty acids (Ox-PUFAs) compared to NS MEFs (Figures 4A-C, S6A, Table S1). Additionally, several lipid classes, including TAGs, PA, PE, PS, and cardiolipin (CL), were among the most downregulated species in senescent MEFs (Figure 4B).

To better understand how α-ESA and α-ESA-me affect the lipidome and drive senolysis, we hypothesized that specific lipid metabolites would be selectively altered in treated S-Eto MEFs but not in NS MEFs. Moreover, if ferroptosis is the underlying mechanism, these alterations should be mitigated by Fer-1 pretreatment. The analysis confirmed that Ox-PUFAs and cholesterol esters (CEs) were upregulated in α-ESA/α-ESA-me treated S-Eto MEFs, but these changes were blunted with Fer-1 pretreatment (Figure 4B). Compared to untreated S-Eto, both α-ESA and α-ESA-me treatment led to a marked increase in individual membrane phospholipids (PC, PE, PI, PS) and cholesterol ester derivative species containing palmitoleic, arachidonate, linoleic, and stearic acid residues (Figures 4D-E). Additionally, there was a notable upregulation of Ox-PUFAs such as arachidonate/linoleic-derived eicosanoids and docosanoids in these treated cells compared to untreated control (Figures 4D-E). However, no significant changes were observed in HNE or related lipid species. Furthermore, pretreatment with Fer-1 reversed many of these changes including CE, CL, PC, PS, and Ox-PUFA, highlighting the ferroptosis-dependence of these lipid changes (Figure 4F). No significant differences in signatures were observed between α-ESA and α-ESA-me, suggesting similar effects (Figure S6B, Table S1).

We also investigated whether there were differences in the abundances of 18:3 species in different experimental groups treated with α-ESAs, which are characterized by their 18:3 structure. Notably, we observed a preferential upregulation of 18:3 fatty acids in TAGs in NS cells, whereas untreated S-Eto cells exhibited elevated levels of 18:3 species in CL and PC forms (Figure S6C). Upon treatment with α-ESAs, S-Eto cells showed a preferential increase in 18:3 species within BMP and CE forms, while NS cells continued to upregulate 18:3 fatty acids in TAGs. Interestingly, Fer-1 pre-treatment reduced the incorporation of 18:3 species into PC, but not CE or BMP forms, in S-Eto cells treated with α-ESAs. Fer-1 had minimal effects in NS cells (Figure S6C). These results suggest that 18:3 fatty acids, likely derived from α-ESAs, are integrated into CEs, CLs, and BMPs, and this integration occurs before ferroptosis is triggered. Additionally, abundance of 18:3 PCs in cells treated with α-ESAs may indicate direct integration of 18:3 α-ESA-containing phospholipids into the cell membranes as an additional mechanism involved in the ferroptotic senolysis.

Taken together, these findings suggest that both α-ESA and α-ESA-me promote the synthesis of arachidonate, stearic and linoleic cholesterol esters, their peroxidation into Ox-PUFAs, and subsequent lipid integration in senescent MEFs, which occur through a ferroptosis-dependent process.

### Structure-based predictions reveal potential targets for α-ESAs in ferroptosis pathways

To identify potential mechanisms through which the α-ESAs induce senolysis thorough ferroptosis, we utilized a deep self-normalizing convolutional neural network (DSNNN) machine learning model to predict compound-pathway interactions. Using KEGG-based classification of human biological pathways, we compared the PathwayMap interaction probabilities of α-ESA and α-ESA-me with a set of known senolytic and ferroptosis-inducing drugs (Figure 5A). The results showed that the two α-ESAs share similar class-wise interaction trends across the panel. Globally, they weakly mimic several known senolytic compounds (e.g., dasatinib, quercetin) and ferroptosis inducers (e.g., erastin and sulfasalazine) in their pathway interaction profiles. However, α-ESA and α-ESA-me share several key classes of pathways with both groups of drugs. They have strong interaction scores in pathways related to amino acid metabolism, replication and repair, cell growth and death, endocrine signaling, aging, cancers, and neurodegenerative diseases. In addition, we used structural features to predict key physicochemical properties of these compounds. The resulted show that, based on structural moieties, both α-ESA and α-ESA-me have high oral bioavailability, blood-brain barrier permeability, as well as low probabilities of systemic toxicity compared to many senolytics and ferroptosis inducers (Figures S7A-C).

**Figure 5.**
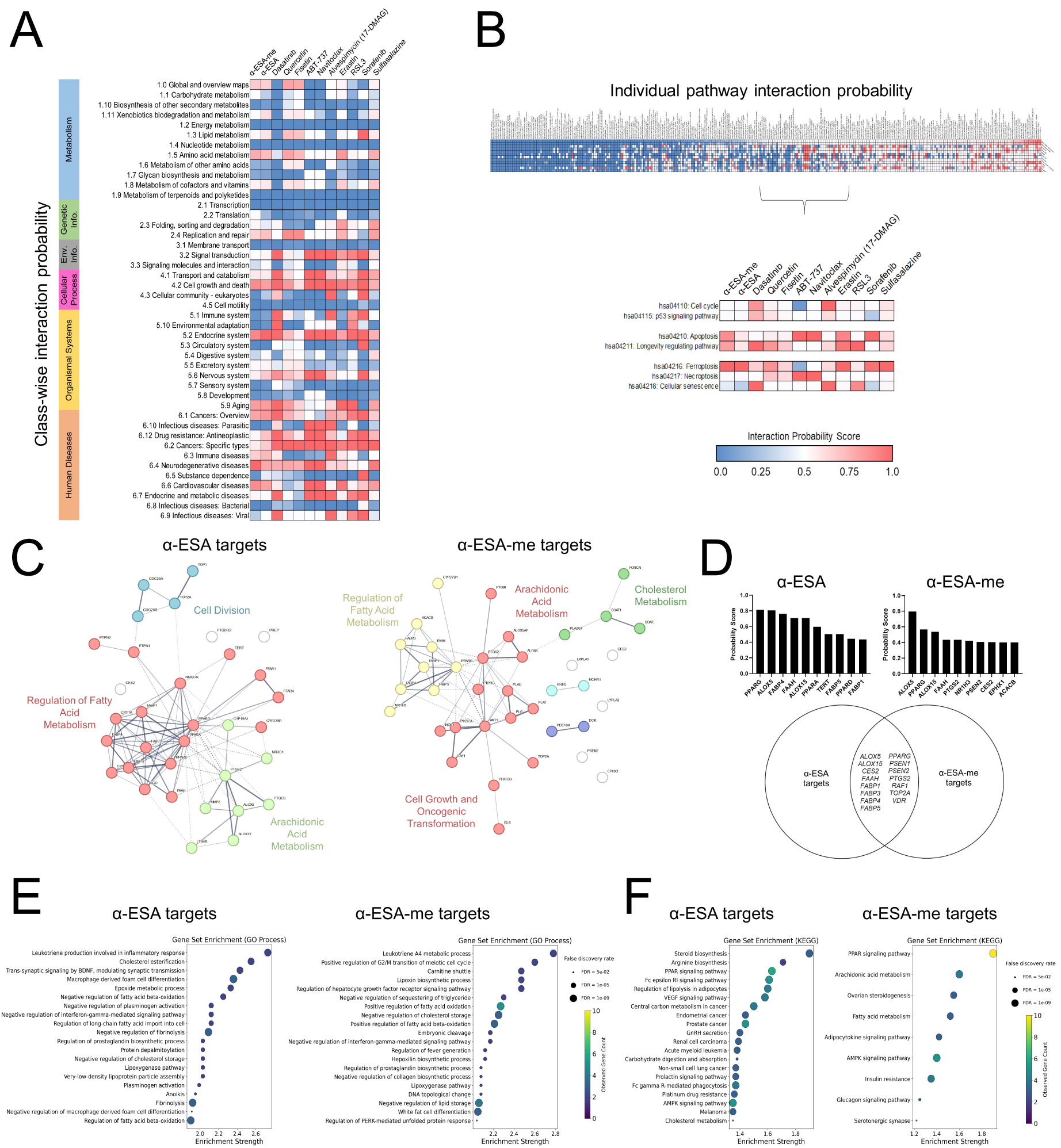
In-silico prediction of pathway interactions and molecular targets for α-ESA and α-ESA-me. (A-B) Class-wise and individual pathway interaction probability scores of compound-pathway pairs for our lipid senolytics, known senolytics, and ferroptosis-inducer standards. (C) k-means clustered STRING protein-protein interaction maps constructed from predicted α-ESA and α-ESA-me target proteins. Labels indicate top GO/KEGG pathway enriched for each cluster. Nodes are clustered based on degree centrality, and edges are defined as level of interaction confidence (probability >0.9). (D) Top 10 structurally probable targets and Venn diagram of overlapping nodes between α-ESA and α-ESA-me interaction networks. (E) GO Process and (F) KEGG Functional enrichment plots for constructed interaction networks of target protein sets.

Further examination of interaction scores with individual KEGG pathways again revealed the top pathways for the α-ESAs to be ferroptosis, cell death, and longevity regulation along with several disease networks (Figure 5B). While both α-ESA and α-ESA-me showed strong propensities for ferroptosis, there was relatively moderate to weak interaction probabilities predicted with apoptosis, necroptosis, cellular senescence, and cell cycle regulation in contrast to the senolytic standards (Figure 5B). Similar trends were seen when examining Reactome pathways (Tables S2B-D). These results suggest that the senolytic activities of the α-ESAs may involve indirect mechanisms, such as ferroptosis induction, rather than directly targeting protein nodes within the cell cycle arrest and senescence signaling networks.

To verify these predictions, we employed the SwissTargetPrediction algorithm for high-throughput screening of ligand-receptor interaction targets based on structural features and ligand similarities of the α-ESAs. The resulting target list was then used to construct clustered protein-protein interaction networks for each α-ESA (Figure 5C, Tables 2E-F). Analysis of network degree centrality revealed overlapping hub genes/proteins between the two networks, including PPARG, PTGS2, and FABP1/3. Expectedly, k-means clustering further identified major clusters within the interaction networks to be predominantly associated with metabolism, oxidation, and the regulation of arachidonic acid, fatty acid, and cholesterol metabolism. Notably, the algorithm identified two arachidonate lipoxygenases (LOXs), ALOX5 and ALOX15, as the most statistically probable targets shared by both α-ESA and α-ESA-me (Figure 5D). Interestingly, ALOX15 and branches from the different arachidonic acid metabolic pathways constitute essential components of the KEGG ferroptosis induction and regulation pathway (https://www.genome.jp/pathway/hsa04216). In accordance with the pathway interaction predictions, neither α-ESA nor α-ESA-me demonstrated strong interaction probabilities with senescence-associated proteins or direct cell cycle regulators other than TOP2A. This is further elucidated through pathway enrichment analysis of the target proteins predicted for each compound. Both α-ESA and α-ESA-me target protein networks showed strong enrichment in pathways related to lipid oxidation, fatty acid metabolism, inflammatory response, and cell death (Figures 5E-F, Tables S2G-H). These predictive algorithms thus support an indirect, ferroptosis-associated interaction model for these two lipid senolytics and their corresponding therapeutic target proteins.

### Transcriptomic analysis further confirms ferroptosis-associated targets

To validate our mechanistic cues in experimental systems, we performed RNA-Seq analysis on non-senescent and senescent WT MEF cells treated with the two lipid senolytics, both in the presence and absence of the ferroptosis inhibitor Fer-1 (Table S4). Gene set enrichment analysis (GSEA) revealed a significant upregulation of genes associated with senescence and SASP in senescent MEF cells compared to non-senescent control (Figure S8). Additionally, there was an enrichment of NF-κB signaling and other inflammatory response pathways in the senescent cells, confirming their senescent phenotype (Figure S8). Upon treatment with α-ESA for 24 hours, we observed the activation of pathways related to reactive oxygen species, heme metabolism, and cholesterol homeostasis (Figure 6A). Pretreatment with the ferroptosis inhibitor Fer-1 significantly altered the transcriptomic response to α-ESA, resulting in the suppression of these pathways (Figure 6B). Similarly, α-ESA-me treatment led to the activation of heme and fatty acid metabolism pathways (Figure 6C), which were also suppressed upon Fer-1 pretreatment (Figure 6D). These findings suggest that both α-ESA and α-ESA-me effectively initiate a ferroptosis-associated transcriptional program and ferroptosis plays a key role in the selective elimination of senescent cells by these compounds.

**Figure 6.**
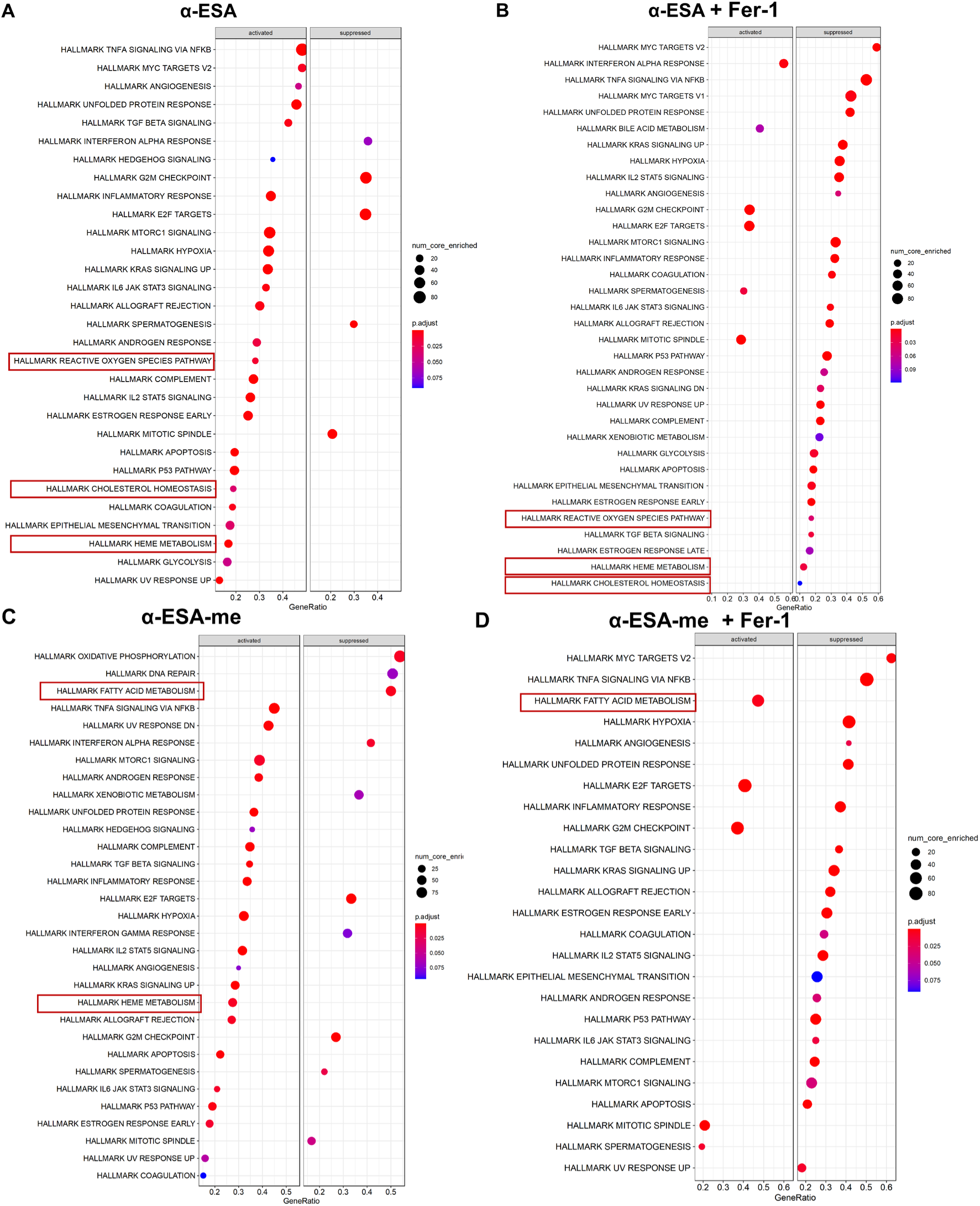
GSEA analysis of α-ESA and α-ESA-me treated senescent cells. Senescent WT MEF cells induced by etoposide were treated with α-ESA (2 µg/mL) and α-ESA-me (2 µg/mL) for 24 hours, with or without 1-h pretreatment of Fer-1 (2 µM) and subjected to RNAseq analysis. Dot plots of enriched pathways determined from GSEA results illustrating hallmarks biological processes associated with (A) α-ESA treatment, (B) α-ESA treatment plus ferroptosis inhibitor Fer-1, (C) α-ESA-me treatment, and (D) α-ESA-me plus ferroptosis inhibitor Fer-1. The gene ratio is defined as the ratio of the count of core enrichment genes to the count of pathway genes. n = 3.

We also examined the mRNA expression levels of our predicted target protein nodes that were shared between α-ESA and α-ESA-me, as well as key ferroptotic proteins. These include the System X_C_-complex (SLC3A2 encoding 4F2, and SLC7A11 encoding xCT) and GPX4 in the cystine-glutathione metabolism arm, as well as ACSL4, LPCAT3, and ALOX15 in the arachidonic acid metabolism arm of the ferroptosis pathway. Among the candidate targets, the most substantial changes were seen in TOP2A, FABP4, and SLC7A11 (downregulated in senescent cells), and ALOX5, ALOX15, and FABP3 (upregulated in senescent cells) (Figure S9A). We verified the expression levels of key ferroptotic nodes using RT-qPCR, where ALOX15 and ALOX5 had the strongest upregulation in senescent cells (Figure S9B). We observed the greatest transcript-level changes for SLC7A11 and PTGS2 when cells are treated with α-ESA and α-ESA-me, but not in non-senescent vs senescent comparisons. These effects are reversed with Fer-1 pretreatment, suggesting these changes are likely dependent on activated ferroptotic processes. Conversely, there were minimal changes in ALOX15 expression in α-ESA/ α-ESA-me treated cells compared to untreated senescent cells.

### α-ESAs are predicted to bind and modulate proteins in the ferroptosis pathways

To evaluate the interactions of the α-ESAs with key proteins in the ferroptosis signaling pathway, we performed flexible molecular docking simulations using a set of enzymes from the KEGG ferroptosis network based on enzymatic reactions contributing to ferroptosis progression and known drug targets. System X_c_- and GPX4 act as important inhibitors or mediators of ferroptosis initiation and progression, where the ACSL4/LPCAT3/ALOX15 axis acts as direct or indirect activators of lipid peroxidation, esterification, membrane integration and ferroptotic processes. A list of receptor crystal structures, native substrates/ligands, cofactors, positive controls used in the docking simulations, and relevant references are provided in supplementary Tables S3A-B.

Biophysical energetic estimations of the ligand-receptor interactions of the α-ESAs showed weak to moderate competition with the native substrates in the cystine/erastin-binding site in the 4F2 heavy chain of the X_C_-complex, and the catalytic reducing site of GPX4; however, these were inconsequential when compared to known inhibitors such as erastin and ML162 (Figure 7A). In case of ACSL4 and LPCAT3, α-ESA but not α-ESA-me show a relatively strong binding affinity towards the substrate catalytic sites outcompeting the native ligands arachidonic acid (ARA) and arachidonoyl-CoA (AR-CoA), although not as strongly as known inhibitor drugs. Looking at ligand-receptor interaction plots, this can be explained by the substrate-receptor and drug-receptor interactions primarily being driven by strong covalent, ionic, and hydrogen bonding events whereas α-ESA and α-ESA-me interactions are chemically preferential towards establishing weak hydrophobic pi-pi and pi-alkyl bonds (Figure S10). Finally, in case of the ALOX15 activation site, both lipid senolytics had substantially stronger binding affinities compared to other selected targets, with α-ESAs outperforming the known ALOX15 allosteric modulator PKUMDL_MH_1001. The 3D ligand-receptor complex with the ALOX15 monomer showed that the binding free energies of the α-ESAs were primarily contributed by hydrogen bonds and a series of weak hydrophobic interactions with amino acid residues in the allosteric activation cavity including PHE88, TRP109, MET148, LEU172, ILE174, LYS175, LEU389, PHE399, ILE403, ARG407, TYR408 (Figures 7B-C). In contrast to other targets, the abundance of these non-polar amino acids in the allosteric activation site in the interdomain cavity neighboring the substrate binding site in ALOX15 constitute a moderately hydrophobic cleft contributing to α-ESA and α-ESA-me binding energetics (Figure 7D). However, lipid molecules, such as arachidonic acid and linoleic acid, are actively bound and oxidized in the more hydrophobic catalytic active site or substrate binding site of ALOX15. If the α-ESAs do not have a preferential selection for binding sites, this can lead to promiscuity in terms of enzyme regulation and possible catalysis of the compounds by ALOX15.

**Figure 7.**
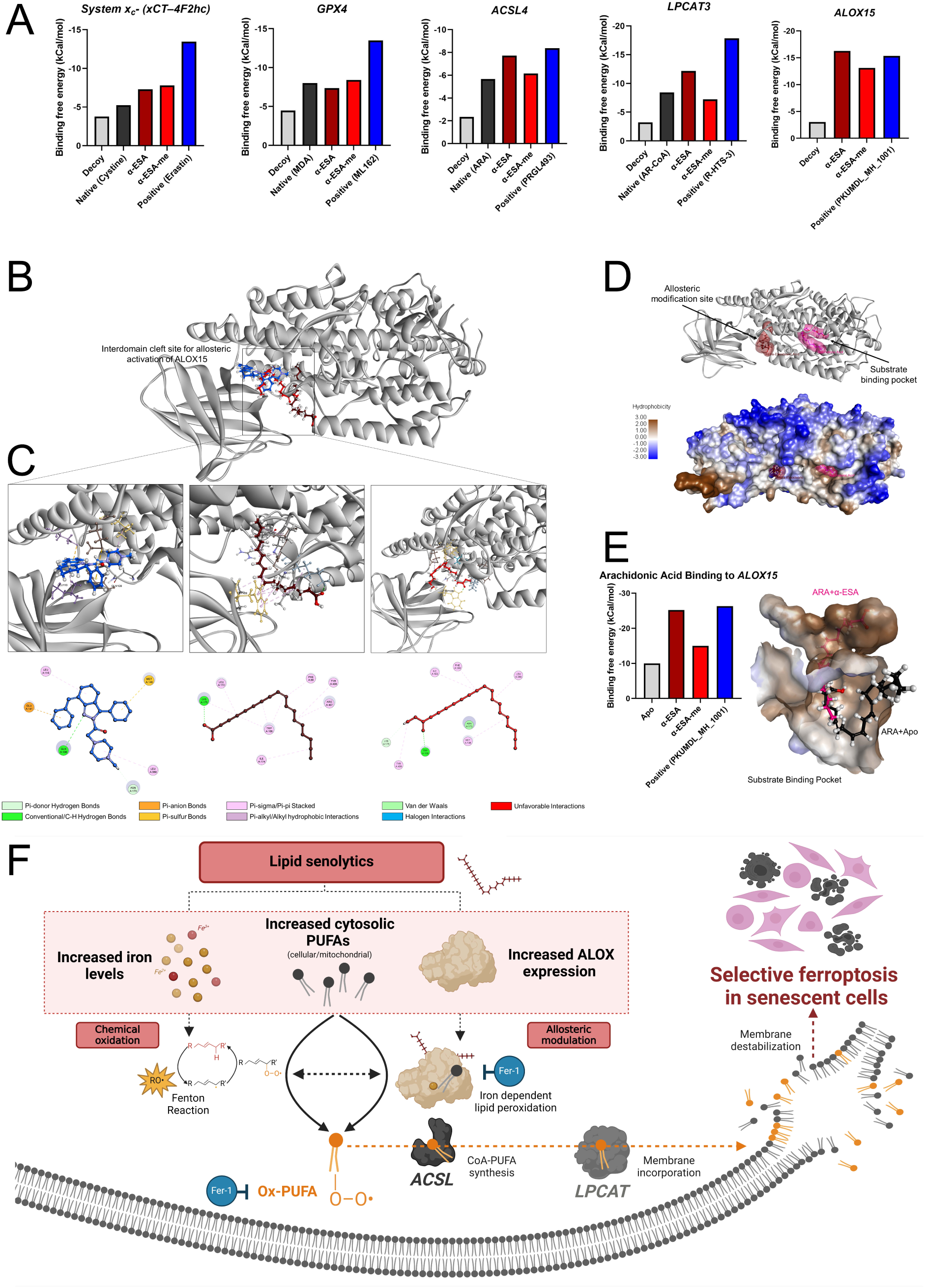
Flexible molecular docking of ferroptosis associated proteins and proposed mechanism. Native substrate or ligand and known inhibitor or activator drug for each protein target was used along with a randomly generated structural decoy/dud molecule. (A) Binding free energy scores of best fitting conformers for each ligand within the selected receptor binding pockets, lower binding free energies correspond to stronger ligand-receptor interactions. Substrate binding pockets for all receptors were predefined except ALOX15 that used the interdomain cleft site for allosteric activation. (B) 3D molecular orientation of best fitting conformers of all simulated ligands in complex within the ALOX15 monomer allosteric activation pocket (color identifiers correspond to panel a). (C) 3D and 2D interaction plots of ligands in the activation pocket with neighboring pocket amino acid residues; ligand-residue interaction ligands are provided under the plots. (D) Protein backbone and hydrophobicity surface projections of ALOX15 monomer with the substrate binding (pink) pocket occupied by native crystalized arachidonic acid (ARA), and allosteric activation (red) pocket occupied by docked α-ESAs. (E) Binding free energy scores of best fitting conformers of ARA in the ALOX15 substrate binding pocket with or without (apo) lipid candidates or drugs docked into the allosteric activation. (F) Proposed mechanism of α-ESA and α-ESA-me induced ferroptotic cell death in senescent cells. Senescent cells exhibit elevated levels of cytosolic PUFAs, cellular ferrous iron and lipoxygenase expression. α-ESA and α-ESA-me treatments exacerbate lipid ROS stress through direct oxidation and/or allosteric activation of ALOX15 activity, promoting the propagation of lipid peroxides. Fer-1 is an anti-ferroptotic agent that directly inhibits ALOX15 and suppresses α-ESA/α-ESA-me mediated Lipid ROS generation. Interactions of reactive lipid species with ACSL and LPCAT enzymes lead to esterification, phospholipid (PL) membrane incorporation and destabilization. *Created in BioRender.com*.

To further investigate this, we performed additional docking experiments for α-ESA and α-ESA-me within a smaller simulation grid in the substrate binding pocket of ALOX15. The results showed that although α-ESA and α-ESA-me had weak to moderate affinity for the substrate binding site (Figure S10), they were not able to competitively inhibit ALOX15-ARA native substrate interactions effectively like the known inhibitor drug Zileuton. The binding energetics, when compared to the allosteric site, also revealed a preferential bias by both lipid senolytics towards the allosteric activation cavity. In all cases, α-ESA showed a stronger binding affinity to most targets compared to α-ESA-me, probably due to the steric hindrance and unfavorable donor-donor interactions with residues in the binding cavities by the additional methyl group on α-ESA-me.

Finally, we inquired whether there are any structural changes to the ALOX15-ARA catalytic binding due to occupancy of the allosteric pocket by α-ESA or α-ESA-me. We performed sequential docking of α-ESAs/PKUMDL_MH_1001 in the allosteric activation site followed by iron cofactor and ARA binding in the substrate binding site of ALOX15. Consistent with previous literature reporting a PKUMDL_MH_1001 mediated increase in arachidonic acid binding by ALOX15,^34^ we found that the binding of the activator drug, as well as both lipid senolytics, increased the binding affinity of ARA in the native substrate binding cavity. Particularly, α-ESA-mediated allosteric distortion of the interdomain cleft in the complex provided better access for the substrate fatty acids into the hydrophobic catalytic cavity of ALOX15 compared to the apo-enzyme (Figure 7E).

## DISCUSSION

Using senescent cell-based phenotypic drug screening assay, here we identified a class of conjugated polyunsaturated fatty acids, α-eleostearic acid (α-ESA) and its methyl ester derivative (α-ESA-me), as novel senolytics. We demonstrated that α-ESA and α-ESA-me effectively kill multiple types of senescent cells induced with different stressors. The senolytic potential of α-ESA and α-ESA-me was further confirmed in naturally aged and accelerated aging mouse models where they significantly reduced senescence in multiple tissues and extended healthspan. Given that clearance of senescent cells mitigates a variety of age-related conditions,^15,16^ these novel lipid senolytics have the potential for the treatment of many age-related diseases driven by senescence.

Most known senolytics induce senescent cell death through apoptosis.^21^ For example, the senolytic combination of dasatinib and quercetin and Bcl-2 family inhibitors, were discovered using bioinformatic screening based on senescent cell anti-apoptotic (SCAP) networks.^35^ In contrast, our mechanistic studies revealed that these lipid senolytics induced senescent cell death through ferroptosis, rather than apoptosis or necrosis. Ferroptosis is an iron-dependent form of cell death characterized by the accumulation of ROS and lipid peroxides through the Fenton reaction.^33^ We demonstrated that senescent cells have elevated levels of ferrous iron and ROS compared to non-senescent cells. The elevated iron overload in senescent cells facilitates the Fenton reaction to produce highly reactive hydroxyl radicals (RO•). α-ESA and α-ESA-me further amplify this process by contributing to the formation of lipid peroxides (PUFA-OO•) and hydroperoxides (PUFA-OOH). These radicals initiate the peroxidation of polyunsaturated fatty acids to form lipid radicals. Interestingly, our lipidomic datasets using etoposide and H_2_O_2_ treated MEFs suggest these senescent cells to be fertile grounds for ferroptosis initiation with upregulated levels of stearic, linoleic, palmitoleic and arachidonate residues hydrolyzed from the membrane. We also showed that 18:3 species were primarily incorporated into cholesterol esters and phospholipids rather than being hydrolyzed into free fatty acids in α-ESA or α-ESA-me treated cells, likely as Ox-PUFAs. Additionally, we observed a preferential incorporation of 18:3 lipids into BMPs and phospholipids in senescent cells, whereas NS MEFs favored their incorporation into TAGs. This difference in lipid storage and incorporation may partially explain the selectivity of α-ESAs in senolysis.

With the accumulation and integration of lipid peroxidation products promoted by the α-ESAs, senescent cells are highly vulnerable to membrane damage which culminates in ferroptotic cell death. The ferroptosis inhibitor Fer-1, acting as a lipid peroxidation inhibitor, can reduce the oxidative stress and lipid peroxidation. The enzyme glutathione peroxidase 4 (GPX4), which is upregulated in senescent cells,^36^ normally mitigates lipid peroxidation by reducing lipid hydroperoxides to their corresponding alcohols (PUFA-OH) using glutathione (GSH) as a cofactor. This also explains why there is no upregulation of Ox-PUFAs and HNEs in untreated senescent MEFs compared to non-senescent ones. However, the high levels of lipid peroxides upon treatment with the α-ESAs likely overwhelm GPX4 activity and overcome ferroptosis resistance in these cells. Our inhibition studies using ferroptosis inhibitors such as ferrostatin-1 and liproxstatin-1 as well as iron chelators like deferoxamine, effectively prevented α-ESA-induced senescent cell death. This confirms the pivotal role of ferroptosis in the senolytic action of α-ESAs. In contrast, inhibitors of apoptosis and necrosis pathways did not offer protection, further validating the unique mechanism of ferroptotic senescent cell death.

Based on our structure-activity relationship study, the ferroptosis-inducing activity of the α-ESAs can be at least partially attributed to their structural feature of conjugated polyunsaturated fatty acids, which are particularly prone to oxidation. In contrast, the double bonds of unconjugated PUFAs are disrupted by a methylene (CH_2_) moiety, thus losing the radical propagation ability. Although many unconjugated PUFAs exhibit various therapeutic functions, such as fish oils (DHA, EPA, and ALA),^26,27^ they showed no senolytic activity. Our SAR analysis also highlights that the configuration of double bonds (*cis*/*trans*) in the carbon chain of fatty acids is a critical determinant of their senolytic activity and selectivity. Additionally, the modification of these polyunsaturated fatty acids with ester groups also significantly influences their senolytic effect. Esterification appears to decrease senolytic activity and but increase selectivity over non-senescent cells. Whether this is due to enhanced cellular uptake or differentiated regulation by certain proteins in the larger senescent cells compared to non-senescent cells remains unknow and warrants further investigation. This SAR information provides a basis for designing more effective lipid senolytics in the future. It is also possible that other conjugated PUFAs may exhibit similar or better senolytic activities than α-ESAs.

Our structure-based DSNN/DCNN machine learning/ligand similarity models and molecular interaction simulations suggest that the α-ESAs represent novel ferroptosis inducers that are safe and bioavailable. They are predicted to exert their senolytic activity through interactions with multiple nodes in the arachidonic acid metabolic arm of the ferroptosis pathway. Recent studies parsing mechanisms underlying ferroptosis have highlighted Fenton chemistry and ALOX activity synergistically contribute to iron-dependent cell death induction and have inter-pathway crosstalk using oxidized metabolites.^37,38^ However, further mechanistic studies are needed to determine whether iron dependent oxidation of these lipid senolytics or enzymatic activation of lipid peroxidases is the primary or exclusive mechanism at work, or if the intermediate components of the interconnected pathways may orchestrate to push senescent cells towards ferroptosis. Multiple studies have shown that the expression of iron-dependent ALOX5/ALOX15, along with their lipid peroxide products, increases during senescence programming.^39–41^ Reinforcing our prediction results, RSL3, a well-known ferroptosis inducer, was shown to selectively clear senescent cells, which was substantially diminished when LOX activity was inhibited by Zileuton.^41^ Indeed, we found that treatment with known ferroptosis inducers, such as the GPX4 inhibitor RSL3 or the glutamate-cysteine antiporter inhibitor erastin, achieves senolysis albeit with low selectivity (Figure S5). Cancer cells, which also accumulate high levels of cellular iron, have similarly shown susceptibility to ferroptotic cell death induced by α-ESA, which was attributed to membrane incorporation via ACSLs and increased lipid hydroperoxides.^42^ Interestingly, these cells had significant enrichment for lipid peroxides with arachidonate residues, suggesting that LOX-mediated PUFA peroxidation may drive ferroptosis in these cells. Our mechanistic study also highlights α-ESAs to be moderately strong interactors in the ACSL4 and LPCAT3 substrate binding cavities, which may explain their observed integration into cholesterol esters, and potential to disrupt phospholipid membranes. Fer-1, an anti-ferroptosis agent, has been reported to exert its effects through direct inhibition of ALOX15 catalytic activity.^43^ We observed a suppression of selective senolysis by α-ESAs in Fer-1 pre-treated cells, suggesting the activity to be at least partially modulated by ALOX15. Our transcriptomic analyses validated a significant upregulation of ALOX15 in senescent cells, and pro-ferroptotic programming upon treatment with the α-ESAs. These effects were reversed with Fer-1 pretreatment, corroborating ferroptosis as a novel mechanism of senescent cell death induced by the α-ESAs.

Iron dyshomeostasis, an abundance of cytosolic lipid peroxidation substrates, and increased ROS levels have been observed in certain senescent cells.^39,40,44–46^ Our findings further demonstrated elevated levels of ferrous iron and ROS, along with the overexpression of ALOX15 in senescent cells compared to non-senescent cells. These factors likely sensitize senescent cells to ferroptosis, making them more vulnerable when exposed to compounds that promote lipid peroxidation. Mechanistically, we propose that the novel lipid senolytics we identified induce senolysis through ferroptosis by exploiting these key vulnerabilities in senescent cells (Figure 7F). Our findings also suggest that inducing ferroptotic senescent cell death could be a novel therapeutic strategy for senolytic interventions. It is foreseeable that repurposing other ferroptosis inducers could potentially lead to the discovery of additional senolytics.

In summary, we identified a novel class of lipid senolytics through a senescent cell-based phenotypic screening of fatty acids. Conjugated polyunsaturated fatty acids, specifically α-ESA and its methyl ester derivative, effectively eliminated a broad range of senescent cells, reduced tissue senescence, and extended healthspan in aged mice. Importantly, these novel lipid senolytics induced senolysis through ferroptosis, rather than apoptosis or necrosis, by exploiting the elevated iron, cytosolic PUFAs, and ROS levels in senescent cells. Our findings also demonstrated the vulnerability of iron-rich, ALOX15-overexpressing senescent cells to drugs that promote lipid peroxide generation and ferroptosis. This study not only expands our understanding of senolytic mechanisms, but also introduces a novel therapeutic strategy for targeted elimination of senescent cells.

## ACKNOWLEDGEMENTS

Funding for this research was provided by NIH grants R01 AG069819 (PDR), P01 AG043376 (PDR, LJN), U19 AG056278 (PDR, LJN), RO1 AG063543 (LJN), P01 AG062413 (LJN, PDR), U54 AG079754 (LJN, PDR), U54 AG076041 (LJN, PDR), and Nathan Shock Center Pilot Grant (LJZ).

## AUTHOR CONTRIBUTIONS

LJZ, LJN, and PDR designed experiments and interpreted the results. LJZ, OE, HY, BH, BZ, AN, KL, WX, AM, EP, SJM, LA, and RO performed experiments. RS and CSP performed computational and bioinformatic analyses. LJZ, RS, CSP, and PDR prepared the original draft of the manuscript. All authors reviewed the manuscript. LJZ and PDR supervised research. LJZ and PDR secured funding.

## Supplemental information

Spreadsheet

**Table S1. Lipidomics data, related to Figure 4**

Spreadsheet

**Table S2. DSNNN and target prediction data, related to Figures 5 and 7**

Spreadsheet

**Table S3. Flexible docking receptor and ligand information, related to Figure 7**

Spreadsheet

**Table S4. GSEA scores, related to Figure 6**

## METHODS

### Resource availability Lead contact

Further information and requests for resources and reagents should be directed to and will be fulfilled by the lead contact, Paul D. Robbins.

### Compounds and reagents

Fatty acids were purchased from Cayman Chemical (Michigan, USA). Hoechst 33342 was purchased from ThermoFisher (H1399). C_12_FDG was purchased from Setareh Biotech (7188). Formaldehyde 32% was purchased from Electron Microscopy Sciences (15714).

### Cells and mice

Primary *Ercc1^-/-^*mouse embryonic fibroblasts (MEFs) and WT MEFs were isolated on embryonic day 12.5-13.5. In brief, mouse embryos were isolated from yolk sac followed by the complete removal of viscera, lung and heart if presented. Embryos were then minced into fine chunks, fed with MEFs medium, cultivated under 3% oxygen to reduce stresses. Cells were split at 1:3 when reaching confluence. MEFs were grown at a 1:1 ratio of Dulbecco’s Modification of Eagles Medium (supplemented with 4.5 g/L glucose and L-glutamine) and Ham’s F10 medium, supplemented with 10% fetal bovine serum, penicillin, streptomycin and non-essential amino acid. To induce oxidative stress-mediated DNA damage, *Ercc1^-/-^* MEFs were switched to 20% oxygen for three passages. WT MEFs were induced senescence by treating with hydrogen peroxide H_2_O_2_ (200 μM) or etoposide (2 μM) for 24 h, followed by 5 days in normal culture media.

Human IMR90 lung fibroblasts were obtained from American Type Culture Collection (ATCC) and cultured in EMEM medium with 10% FBS and pen/strep antibiotics. To induce senescence, cells were treated with 20 μM etoposide for 24 h, followed by five days in normal culture media.

Human umbilical vein endothelial cells (HUVECs) were obtained from ATCC and cultured using Endothelial Cell Growth Media plus supplement (without vascular endothelial growth factor (VEGF)) and 1% pen/strep antibiotics. The cells were experimentally treated at late passages 13 to 15.

*Ercc1*^+/−^ and *Ercc1*^+/^*^Δ^* mice from C57BL/6J and FVB/n backgrounds were crossed to generate *Ercc1*^−/^*^Δ^* mice to prevent potential strain-specific pathology. Aged wild-type C57BL/6J:FVB/NJ mice were generated by crossing C57BL/6J and FVB/n inbred mice purchased from Jackson Laboratory. Mice were left to age for two years before being enrolled into the late life intervention study. Animal protocols used in this study were approved by the University of Minnesota Institutional Animal Care and Use Committees.

### Method details Senotherapeutic screening

Senescence was evaluated based on SA-β-gal activity using C_12_FDG staining assay. Specifically, senescent *Ercc1^−/−^* MEFs were passaged for three times at 20% O_2_ to induce senescence then seeded at 3000 cells per well in black wall, clear bottom 96 well plates at least 16 hours prior to treatment. Following the addition of drugs, the MEFs were incubated for 48 hours at 20% O_2_. After removing the medium, cells were incubated in 100 nM Bafilomycin A1 in culture medium for 60 min to induce lysosomal alkalinization, followed by incubation with 20 μM fluorogenic substrate C_12_FDG (7188, Setareh Biotech, USA) for 2 h and counterstaining with 2 μg/ml Hoechst 33342 (H1399, Thermo Fisher Scientific, MA, USA) for 15 min. Subsequently, cells were washed with PBS and fixed in 2% paraformaldehyde for 15 min. Finally, cells were imaged with 6 fields per well using a high content fluorescent image acquisition and analysis platform Cytation 1 (BioTek, VT, USA). Cells were seeded at 3000 cells per well in black-wall, clear-bottom 96-well plates at least 16 hours prior to treatment. Following the addition of drugs, cells were incubated for 48 hours. After removing the medium, cells were incubated in 100 nM Bafilomycin A1 in culture medium for 1 hour to induce lysosomal alkalinization, followed by incubation with 20 μM fluorogenic substrate C_12_FDG (7188, Setareh Biotech, OR, USA) for 2 hours and counterstaining with 2 μg/mL Hoechst 33342 (H1399, Thermo Fisher Scientific, MA, USA) for 15 minutes. Subsequently, cells were washed with PBS and fixed in 2% paraformaldehyde for 15 minutes. Finally, cells were imaged with six fields per well using the Cytation 1 Cell Imaging Multi-Mode Reader (BioTek Instruments, VT, USA).

### Cellular ferrous iron measurement

Cells were treated with α-ESA (10 μg/mL) or α-ESA-me (20 μg/mL) for 6 hours with or without a 1-hour pretreatment with Fer-1 (2 μM). After treatment, cells were stained with 1 μM FerroOrange (SCT210, Sigma-Aldrich, USA) for 1 hour and 2 μg/mL Hoechst 33342 (H1399, Thermo Fisher Scientific, MA, USA) for 15 minutes. The staining solution was removed, and cells were washed with PBS. Cells were immediately imaged using the Cytation 1 Cell Imaging Multi-Mode Reader (BioTek Instruments, VT, USA).

### Cellular reactive oxygen species detection

Cells were treated with α-ESA (10 μg/mL) or α-ESA-me (20 μg/mL) for 6 hours with or without a 1-hour pretreatment with Fer-1 (2 μM). After treatment, cells were stained with 1 μM H2DCFDA (D399, Thermo Fisher Scientific, MA, USA) for 1 hour and 2 μg/mL Hoechst 33342 (H1399, Thermo Fisher Scientific, MA, USA) for 15 minutes. The staining solution was removed, and cells were washed once with PBS. Cells were immediately imaged using the Cytation 1 Cell Imaging Multi-Mode Reader (BioTek Instruments, VT, USA).

### Lipid peroxidation assay

Cells were treated with α-ESA (10 μg/mL) or α-ESA-me (20 μg/mL) for 6 hours with or without a 1-hour pretreatment with Fer-1 (2 μM). Following treatment, cells were stained with 10 μM C11 BODIPY (D3861, Thermo Fisher Scientific, MA, USA) for 30 minutes and 2 μg/mL Hoechst 33342 (H1399, Thermo Fisher Scientific, MA, USA) for 15 minutes. The staining solution was removed, and cells were washed with PBS. Cells were immediately imaged using the Cytation 1 Cell Imaging Multi-Mode Reader (BioTek Instruments, VT, USA).

### Healthspan evaluation of *Ercc1^-/Δ^* mice

Healthspan assessment of *Ercc1^-/Δ^*mice was conducted twice per week to evaluate age-related symptoms, including body weight, tremor, forelimb grip strength, kyphosis, hindlimb paralysis, gait disorder, dystonia and ataxia. Kyphosis, body condition and coat condition were used to reflect general health conditions. Ataxia, dystonia, gait disorder and tremor were used as indicators of aging-related neurodegeneration. Muscle wasting was studies by monitoring hindlimb paralysis and forelimb grip strength. All aging symptoms were scored based on a scale of 0, 0.5 and 1, with the exception of dystonia that has a scale from 0 to 5. The sum of aging scores of each group was used to determine the overall aging conditions, with zero means no symptom presented.

### RT-qPCR analysis

Total RNA was extracted from cells or snap frozen tissues using Trizol reagent (Thermo Fisher, USA). cDNA was synthesized using High-Capacity cDNA Reverse Transcription Kit (Thermo Fisher, USA). Quantitative PCR reactions were performed with PowerUp™ SYBR™ Green Master Mix (ThermoFisher, USA). The experiments were performed according to the manufacturer’s instructions. The sequences of the primers used were listed below.

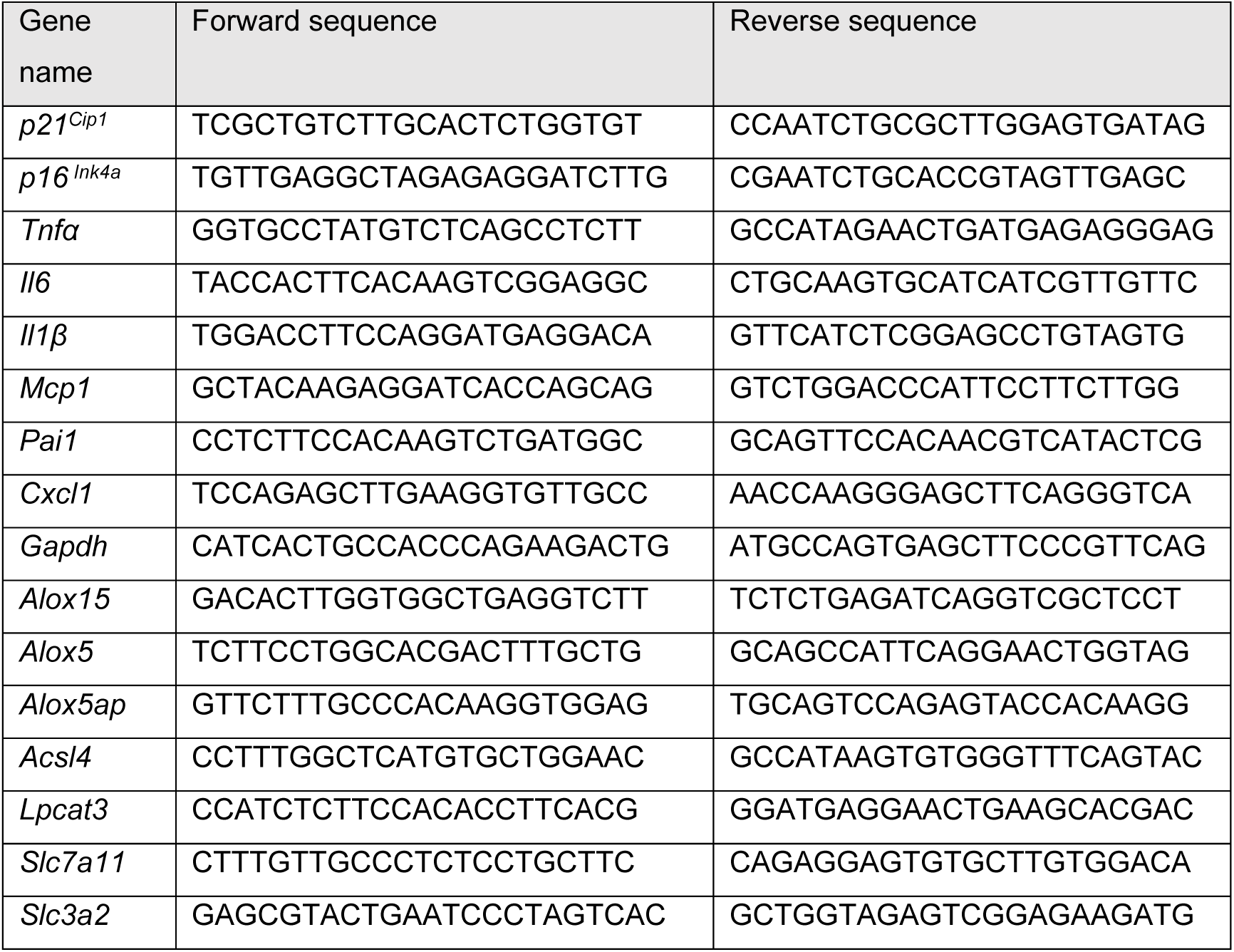
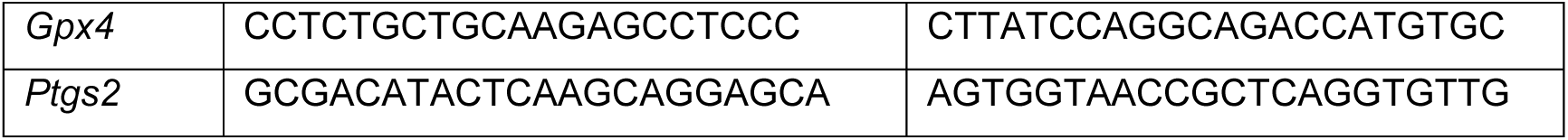

### Lipidomics analysis

Non-senescent and senescent MEF cells were treated with α-ESA (2 µg/mL) or α-ESA-me (2 µg/mL) for 8 hours with or without 1-hour pretreatment with Fer-1 (2 μM). Upon treatment, cells were washed with PBS and about 3×10^6^ cells were collected using trypsin. Cell pellets were homogenized on ice with 0.5 ml of 0.1× PBS at 6500 rpm, 0 °C in 2 ml Precellys Lysing tube (Bertin) using Cryolys Evolution homogenizer (Precellys Evolution). The protein concentration of cell homogenates was determined using the Pierce BCA protein assay kit (#23225, Thermo Scientific). Bovine serum albumin was used as standard. An adequate amount of each homogenate (equivalent to 0.32 mg protein) was transferred into a disposable glass culture test tube. The lipid internal standard mixture for quantitation of lipids was added prior to lipid extraction. Lipid extraction was performed using a modified Bligh and Dyer procedure, as described previously^47^ Lipid extracts were flushed with nitrogen, capped, and stored at -20 °C until lipid analysis.

Shotgun lipidomics was performed as described previously.^48^ Lipid extract was further diluted to a final concentration of ∼500 fmol total lipids per μL. Mass spectrometric analysis was performed on a triple quadrupole mass spectrometer (TSQ Altis, Thermo Fisher Scientific) and a Q Exactive mass spectrometer (Thermo Scientific, San Jose, CA), both of which were equipped with an automated nanospray device (TriVersa NanoMate, Advion Bioscience Ltd, Ithaca, NY) as described.^49^ Identification and quantification of lipid species were performed using an automated software program.^50,51^ Data processing (e.g., ion peak selection, baseline correction, data transfer, peak intensity comparison, and quantitation) was performed as described.^51^ Data were normalized per million cells. Reproducibility by analyzing a prepared sample by multidimensional mass spectrometry-based shotgun lipidomics (MDMS-SL) is approximately 95%, and precision is approximately 90%, largely due to variations in determining protein content for normalizing lipid levels.^49^ The content of lipids was analyzed and quantified by the GraphPad Prism 10 software. Z-Scores, Log2FC in abundance, and p-values were calculated using the pandas, scipy.stats and numpy modules on the Spyder 6 standalone IDE running on Python 3.12. Clustered heatmaps and volcano plots were generated using the pandas, seaborn, matplotlib and scipy packages.

### Molecular pathway interaction probability analysis

Self-normalizing neural-network (DSNNN) based metabolic pathway interaction probability predictions were simulated using the PathwayMap algorithm as previously described.^52^ Canonical SMILES strings for each compound were retrieved from https://pubchem.ncbi.nlm.nih.gov and submitted as inputs (listed in Table a in Source Data 2). Both class-wise and individual pathway results for KEGG and Reactome interaction databases were generated and visualized.

### Pharmacokinetic properties and *in-silico* toxicity screening of compounds

Canonical SMILES of the compounds were subjected to *in-silico* ADME prediction using the SWISSADME web server.^53^ We subsequently performed prediction algorithms for human oral toxicity parameters including mutagenicity, tumorigenicity, allergenicity, reproductive teratogenicity and cytotoxicity of the compounds using the OSIRIS Property Explorer v1.1 standalone software (https://www.organic-chemistry.org/prog/peo) and the ProTox-3.0 prediction tool.^54^

### Structure-based virtual target prediction and network construction

The canonical SMILES strings for the ligands were used as inputs for the SwissTargetPred algorithm^55^. Both 2D and 3D similarity measures were used as search parameters with *H. sapiens* selected as the target database. Probability score of >0.2 was used as statistical cutoff for identified targets and the protein identifiers were retrieved. The set of identifiers for each compound was used as node inputs in the STRING interaction database (https://string-db.org). Results were restricted to *H. sapiens* and submitted list of proteins only. Molecular interaction modes (edges) with highest confidence score (>0.90) were generated and used to analyze highly significant (FDR<0.01) functional enrichment of KEGG pathways and biological processes (gene ontology). Subsequently, nodes and edges were imported to CytoScape v3.9.0 for visualization using the stringApp 1.7.0 add-on.^56^ Node values were assigned using weighted k-shell distribution via wk-shell-decomposition 1.1.0. Subsequently, nodes were clustered using clusterMaker2 2.0 taking undirected edges and wkshell values as array input with granularity of 2.5.

### Ligand and receptor crystal structure acquisition and 3D structure preparation

The 3D ligand structures of candidate compounds and control were retrieved from https://pubchem.ncbi.nlm.nih.gov. 20 decoy molecules per ligand were generated using the DUD-E database (https://dude.docking.org). Prior to docking runs, we optimized the ligand structures through minimization and polar protonation (pH = 7.4). 3D crystal structures of inhibitor-bound/activator-bound open conformations of the target receptors from the RCSB-PDB database (Table S2). The AlphaFold predicted structure of ACSL4 was retrieved from https://www.rcsb.org/structure/AF_AFO60488F1. The structures were optimized using Biovia Discovery Studio Modeling Environment 2024. The crystals were titrated and protonated at pH = 7.4. Co-crystal water, non-cofactor ligand molecules were removed and the appropriate binding cavities were determined by the deep convolutional neural network (DCNN) based protein-binding site prediction algorithm DeepSite^57^ (Score≥9.0), and verified with existing literature. All crystal structures were subjected to the PROCHECK algorithm for stereochemical quality assessment in order to ensure accurate docking pose predictions (https://www.ebi.ac.uk/thornton-srv/software/PROCHECK). Estimations of the whole-model reliability of the 3D structures was performed by evaluating the QMEAN Z-scores of the PDB structures using the ProSA server (https://prosa.services.came.sbg.ac.at/prosa.php). All structures were within acceptable resolution, and Z-scores for NMR and X-ray quality crystal references (Table S2).

### Flexible molecular docking simulation

Structure-based flexible molecular docking of the ligands into the receptor binding cavities were done using the DockThor v2.0 platform.^58^ A receptor search grid of 25Å×25Å×25Å was considered near the binding site of each target protein and post-docking OPLS forcefield minimization was carried out. Monomeric subunits of the proteins were used as receptor files and grids were manually defined around the binding cavity for running the docking simulations. DMRTS method was employed for the simulation with 100,000 evaluations per run with an initial population of 750 and 25 runs per ligand. In addition to lipid and drug molecules, 20 decoy molecules per complex were used to assess the specificity of the docking protocol and the best binding decoy was used for comparisons. Ligand-receptor interactions were scored using the rDock master scoring function, and binding free energy was calculated as *Binding Free Energy (ΔG)= intermolecular+ ligand intramolecular+site intramolecular+ external restraints* (https://rxdock.gitlab.io/documentation/devel/html/reference-guide/scoring-functions.html)k.inPgovsits-duoaclization and ligand-receptor interactions were analyzed using Biovia Discovery Studio Modeling Environment 2024 platform.

### RNA-Seq and enrichment analysis

Non-senescent and senescent MEF cells were treated with α-ESA (2 µg/mL) or α-ESA- me (2 µg/mL) for 24 hours with or without 1-hour pretreatment with Fer-1 (2 μM). RNA samples were extracted using Trizol reagent (Thermo Fisher, USA). Samples were quantified using fluorimetry (RiboGreen assay) and RNA integrity was assessed using capillary electrophoresis, generating an RNA Integrity Number (RIN). All samples had at least 500ng mass and RIN of at least 8. Library preparation was carried out using the Illumina TruSeq Stranded Total RNA Library Prep kit, followed by sequencing on the NovaSeq 6000 using 150 PE flow cell, with a sequencing depth of 20 million reads per sample. Quality control on raw sequence data for each sample was performed with FastQC. Read mapping was performed via Hisat2 (v2.1.0) using the mouse genome (GRCm38.94) as reference. Gene quantification was done via Feature Counts for raw read counts. Data have been deposited in the Gene Expression Omnibus under accession number GSE278372. Differentially expressed genes (DEGs) were identified using the R package edgeR. Gene Set Enrichment Analysis (GSEA) and over representation analysis (ORA) was performed using the R Clusterprofiler package.^59,60^ DEGs with adjusted p-values < 0.05 and log_2_FC > |1| were selected for downstream representation analysis.

### Statistical analysis

Data were statistically analyzed by Graphpad Prism software. Two-tailed Student’s *t*-test was performed to determine differences between two groups, and one-way ANOVA with Tukey’s test was used for three groups. A value of *p* <0.05 was considered as statistically significant, shown as \**p* <0.05, \*\**p* <0.01, \*\*\**p* < 0.001 and ****p<0.0001.

---

**Figure S1.**
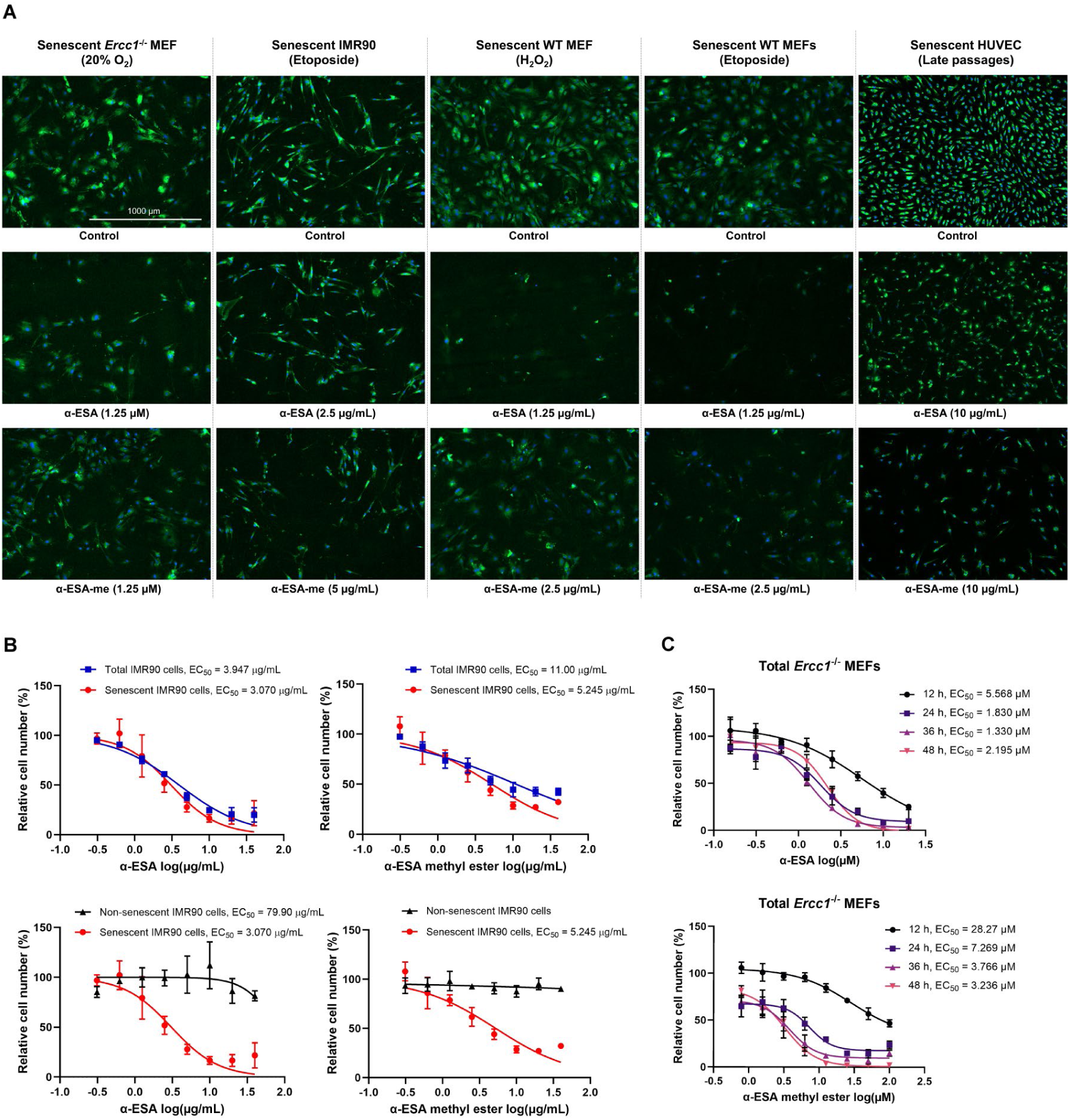
Senolytic activity of α-ESA and α-ESA-me in various senescent cell models, related to Figure 1. (A) Representative images from the C_12_FDG-based SA-β-gal senescence assay in senescent *Ercc1*^-/-^ MEF cells treated with α-ESA and α-ESA-me for 48 hours. Images were captured using Cytation 1 at 4X magnification. Blue fluorescence indicates Hoechst 33324-stained nuclei and green fluorescence marks C_12_FDG-stained SA-β-gal positive senescent cells. (B) Dose-response curves of α-ESA and α-ESA-me in non-senescent and senescent IMR90 cells. Error bars represent SD for n = 3. (C) Time-dependent dose response of senescent *Ercc1*^-/-^ MEF cells treated with α-ESA or α-ESA-me. Error bars represent SD for n = 3.

**Figure S2.**
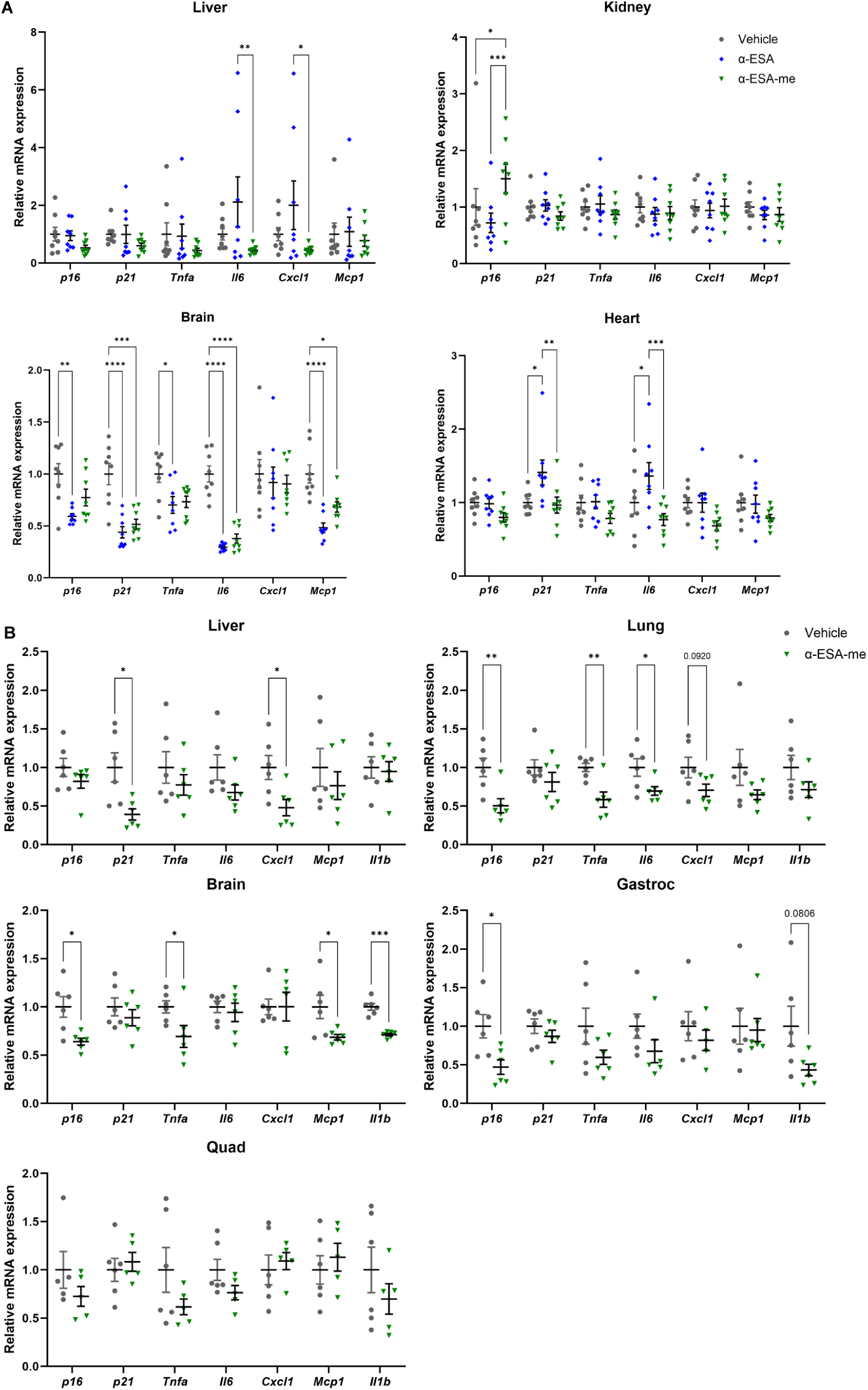
Senolytic effects of α-ESAs in aged WT C57BL/6 mice, related to Figure 2. (A) The effects of α-ESA and α-ESA-me on reducing senescence markers across different tissues in WT C57BL/6 mice aged 20-22 months. Error bars represent SEM for n = 8. (B) The effects of α-ESA-me on reducing senescence across different tissues in WT C57BL/6 mice aged 32 months. Error bars represent SEM for n = 6.

**Figure S3.**
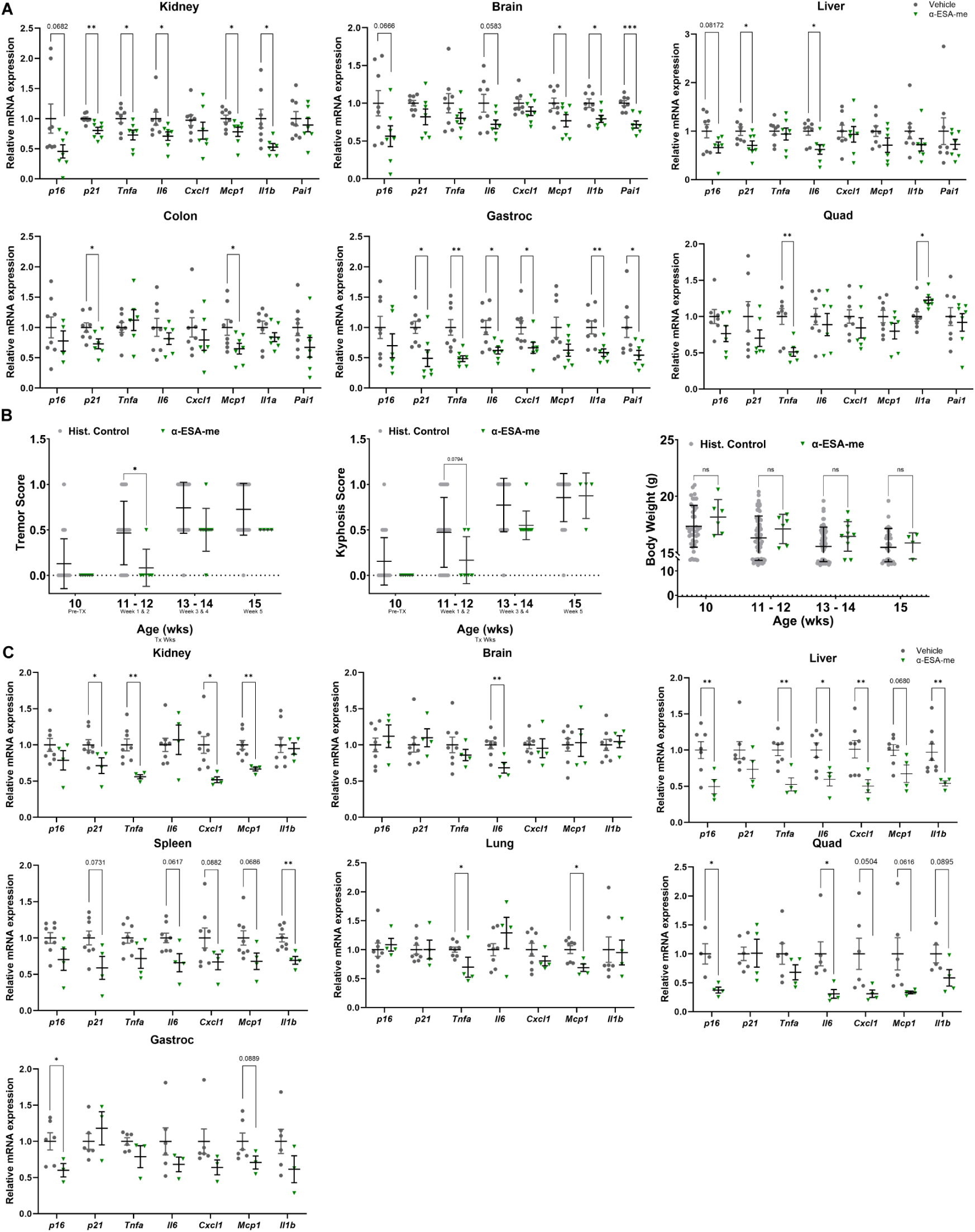
Senolytic effects of α-ESA-me in *Ercc1*^-/Δ^ progeria mice, related to Figure 2. (A) Acute treatment with α-ESA-me reduced tissue senescence in *Ercc1*^-/Δ^ progeria mice. Error bars represent SEM for n = 8 (vehicle) and n = 7 (α-ESA-me). (B) Chronic treatment with α-ESA-me improved aging symptoms in *Ercc1*^-/Δ^ progeria mice. (C) Chronic treatment with α-ESA-me also reduced senescence and SASP markers across multiple tissues in *Ercc1*^-/Δ^ progeria mice. Error bars represent SEM for n = 6 (vehicle) and n = 4 (α-ESA-me).

**Figure S4.**
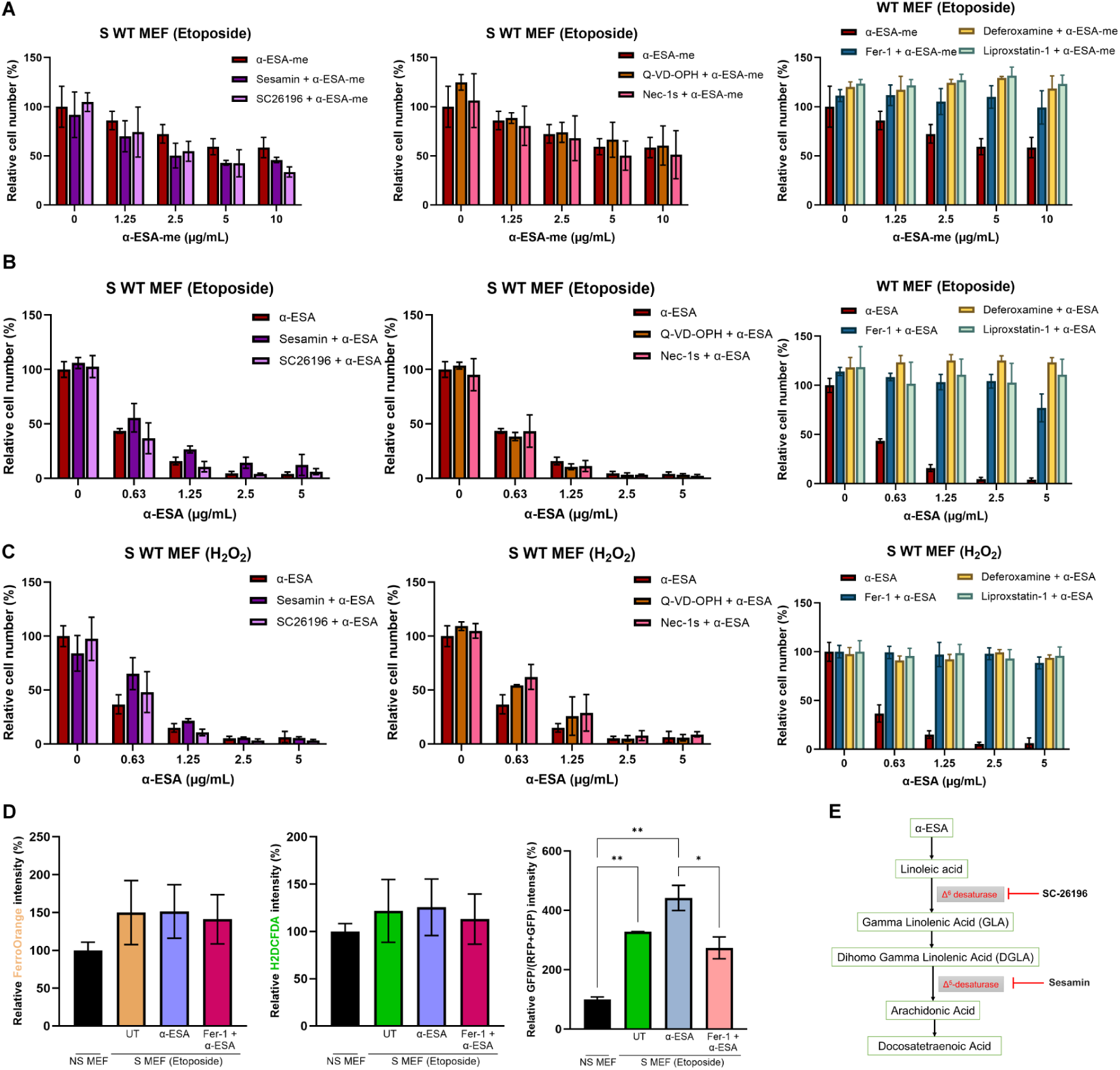
α-ESAs induced senolysis through ferroptosis, related to Figure 3. (A-C) Senescent MEF cells were treated with α-ESA or α-ESA-me for 48 hours, with or without 1-hour pretreatment with the following compounds: sesamin (50 μM), SC26196 (200 nM), Q-VD-OPH (20 μM), Nec-1s (50 μM), Fer-1 (2 μM), deferoxamine (50 μM), or liproxstatin-1 (2 μM). Error bars represent SD for n = 3. (D) Cells were treated with α-ESA-me (5 µg/mL) for 6 hours, with or without 1-hour pretreatment with Fer-1 (2 μM). Ferrous iron, ROS, and lipid peroxidation were detected by FerroOrange, H2DCFDA, and C11 BODIPY, respectively. Error bars represent SD for n = 2. (E) The metabolism of α-ESA to linoleic acid and downstream metabolites.

**Figure S5.**
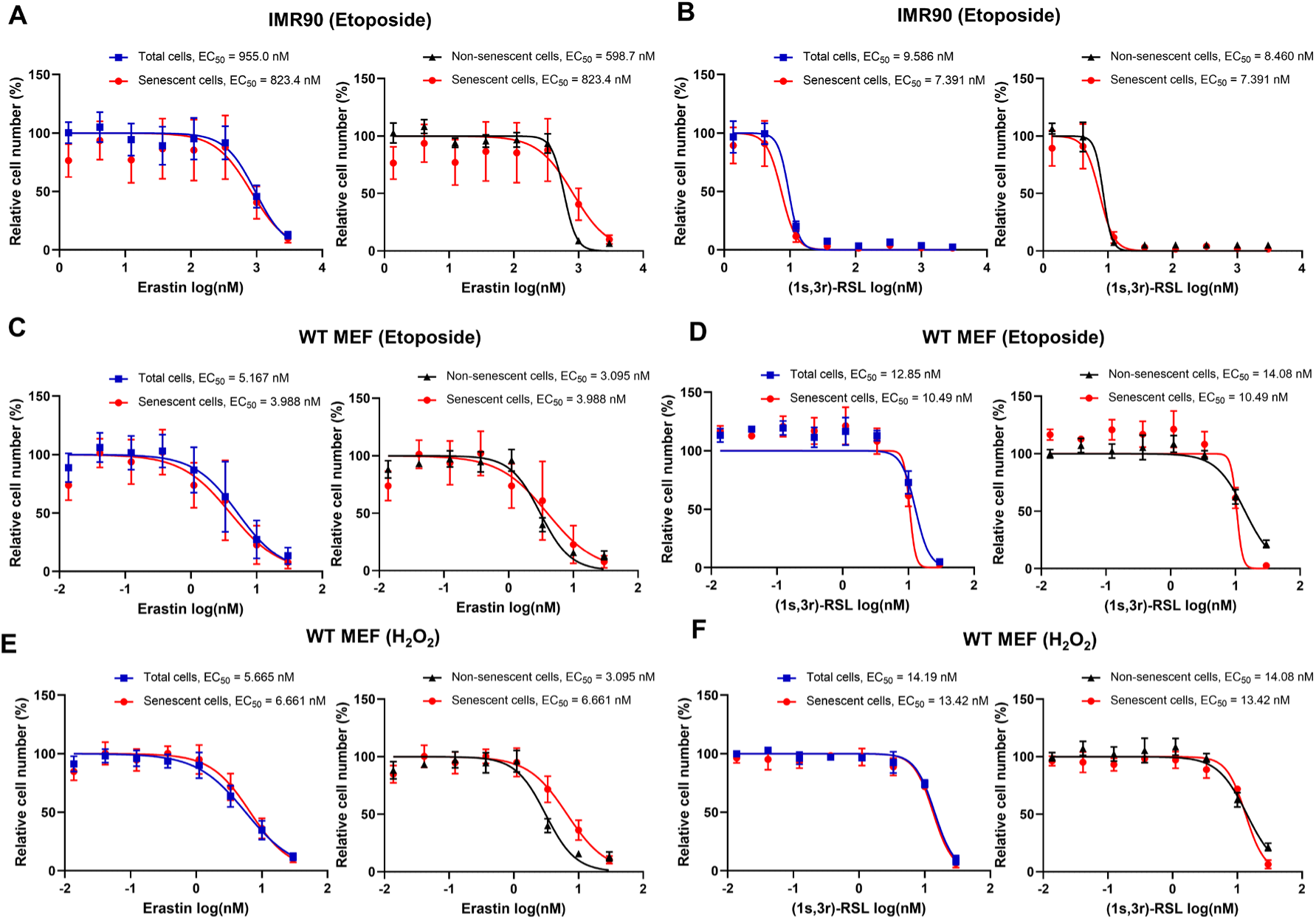
Senolytic effects of ferroptosis inducers, related to Figure 3. Dose-response curves for EC_50_ determination of erastin and (1s, 3r)-RSL in (A, B) senescent IMR90 cells, (C, D) senescent WT MEF cells induced by etoposide, and (E, F) senescent WT MEF cells induced by H_2_O_2_. Error bars represent SD for n = 3.

**Figure S6.**
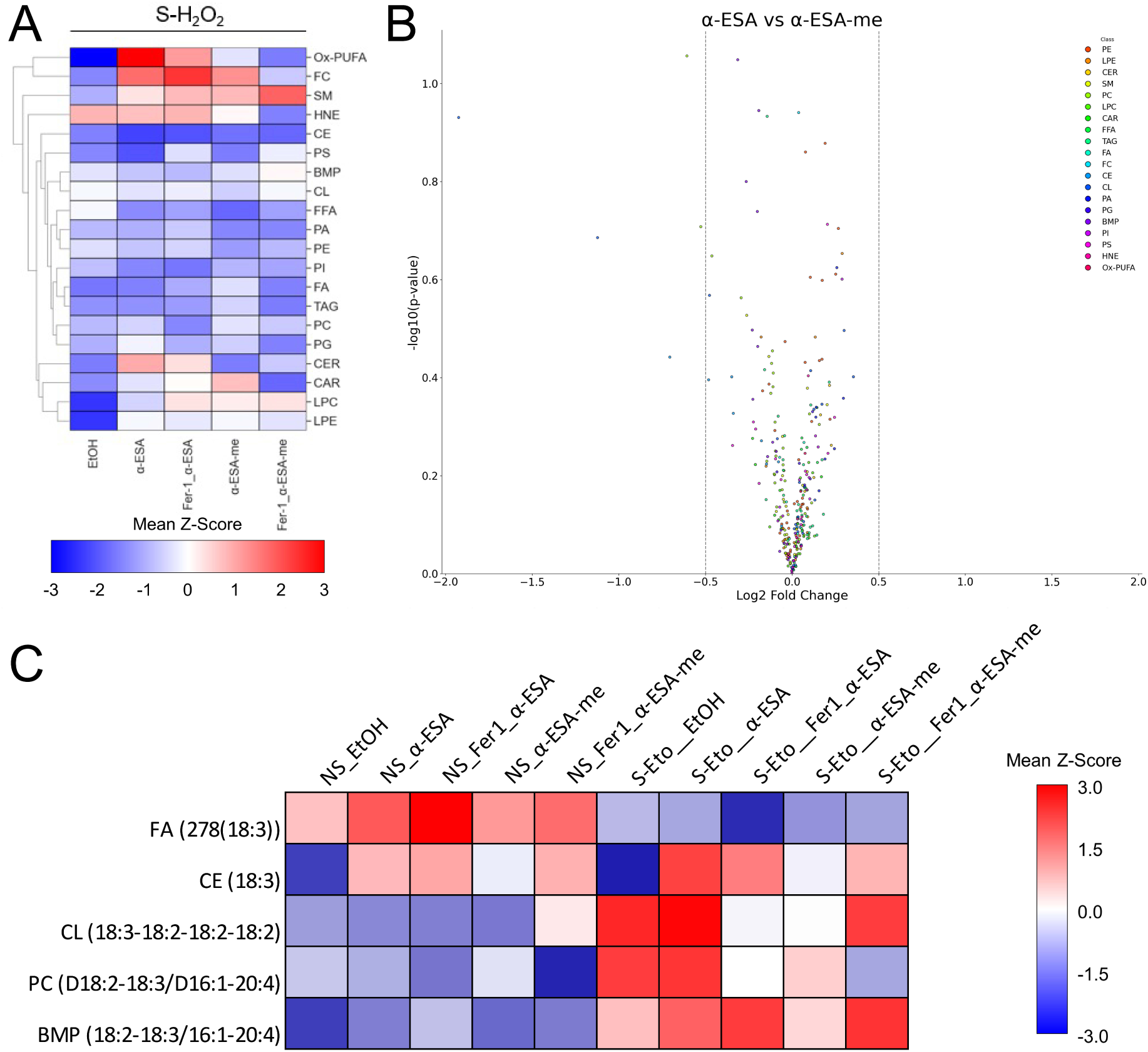
Lipidomic analysis data, related to Figure 4. (A) Euclidian clustered heatmap showing color-coded Z-scores for lipid class-wise distribution in S-H_2_O_2_ MEFs. Data are based on mean Z-scores for species within each class and n= 4 for all groups. (B) Volcano plot of differential changes in abundance of lipid species for α-ESA vs α-ESA-me treated MEFs; colors represent lipid classes. (F) Heatmap of species-wise Z-scores of 18:3 species in different treatment groups, data are based on mean Z-scores for each species and n= 4 for all groups. Abbreviations: PE, Phosphatidylethanolamine; LPE, Lyso-Phosphatidylethanolamine; CER, Ceramide; CE, Cholesterol Esters; SM, Sphingomyelin; PC, Phosphatidylcholine; LPC, Lyso-Phosphatidylcholine; CAR, Acyl-Carnitine; FFA, Free Fatty Acid; TAG, Triacylglycerol; FA, Fatty Acyl Chains in TAG; CL, Cardiolipin; PA, Phosphatidic acid; PG, Phosphatidylglycerol; BMP, Bis(Monoacylglycero) Phosphate; PI, Phosphatidylinositol; PS, Phosphatidylserine; Ox-PUFA, Oxidized Polyunsaturated Fatty Acids.

**Figure S7.**
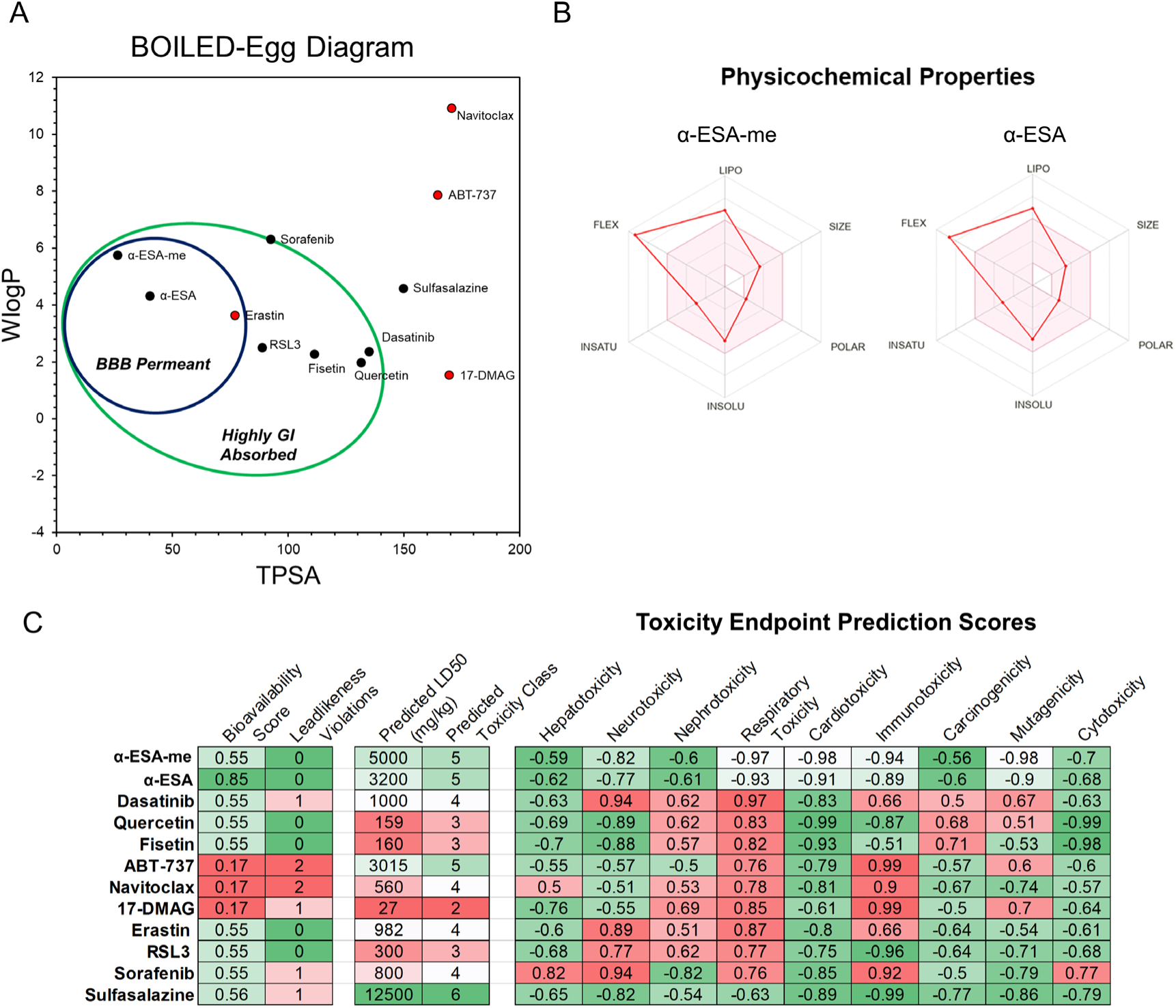
Physicochemical properties of α-ESA and α-ESA-me, related to Figure 5. (A) BOILED-Egg summary diagram of the α-ESAs and standard compounds. The hydrophobicity (WLogP) and topological polar surface area (TPSA) values were used to determine the potential brain-blood barrier (BBB) permeability, GI absorption capacities, and bioavailability scores of the compounds. P-gp substrate molecules have a higher probability of overcoming drug resistance. (B) Physicochemical property maps show high flexibility and lipophilicity for α-ESA and α-ESA-me. (C) Predicted oral bioavailability, drug-likeness scores, and toxicity profiles of the α-ESAs and standard compounds.

**Figure S8.**
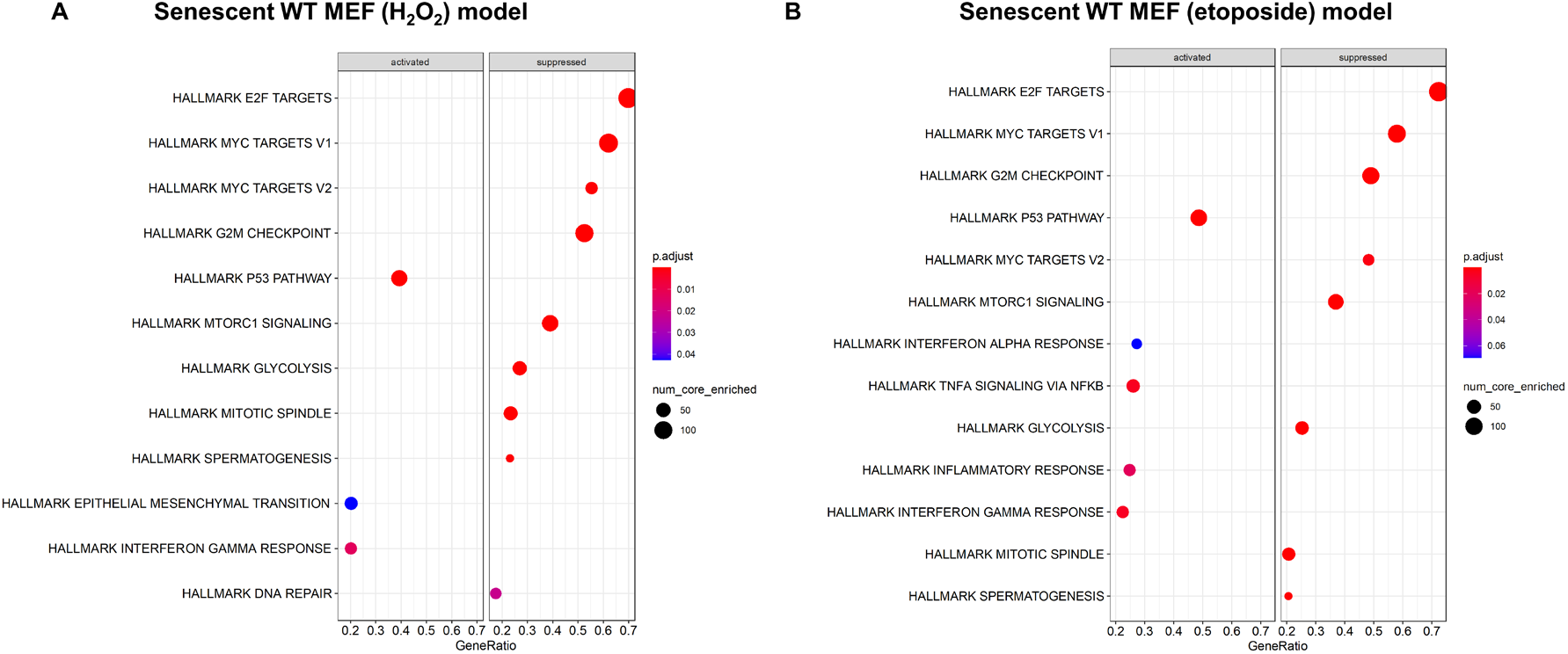
Pathway analysis of senescent cells, related to Figure 6. Dot plots of enriched pathways (GSEA) associated with senescent cells using Hallmark database. (A) WT MEF hydrogen peroxide senescence model (B) WT MEF etoposide senescence model. Hallmark p53 and inflammatory responses pathways are activated in the senescent cells of both corresponding models.

**Figure S9.**
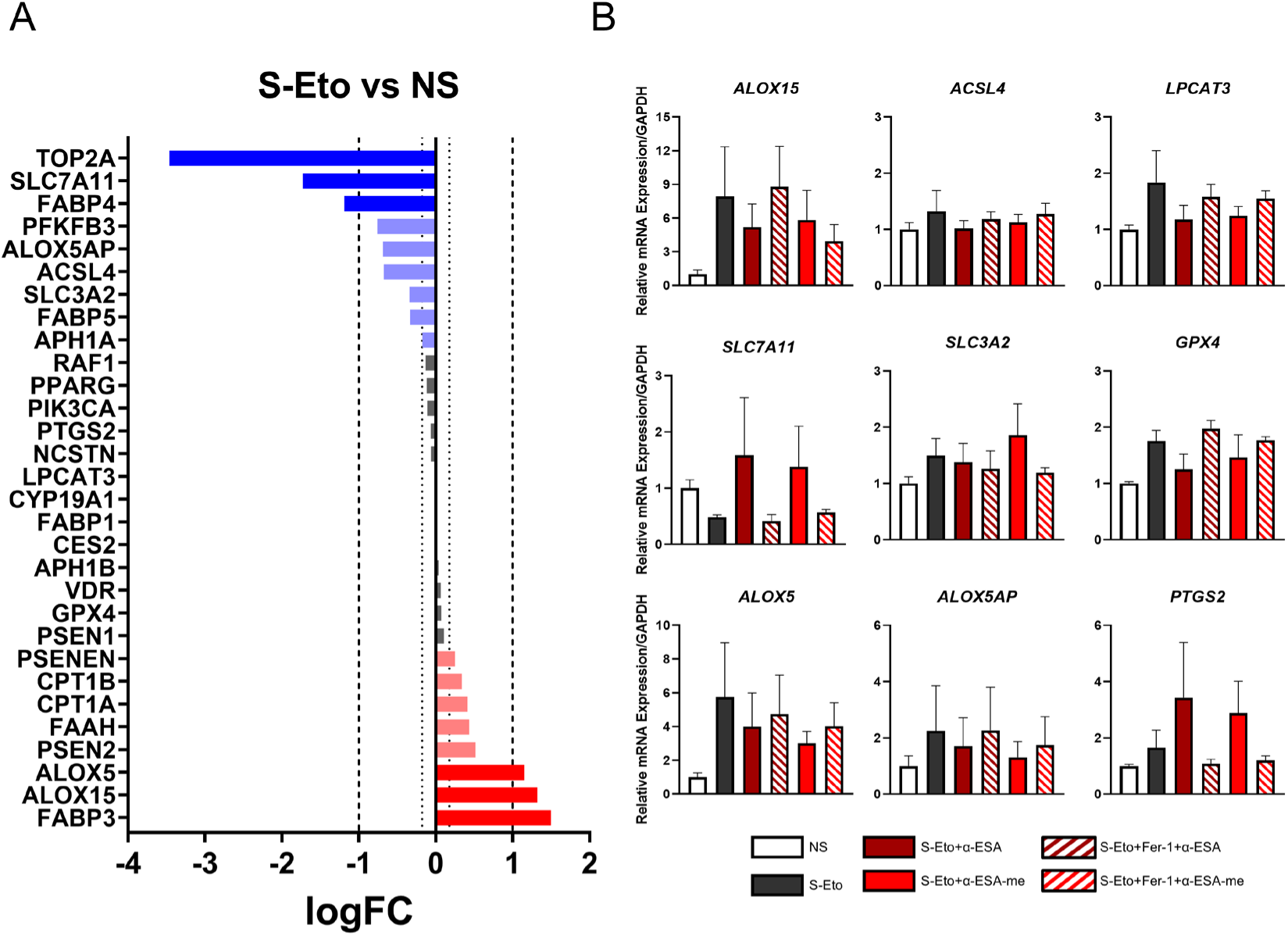
Expression profiles of predicted targets and ferroptosis pathway genes, related to Figure 6 (A) Ranked expression change in etoposide-treated senescent (S-Eto) vs non-senescent (NS) MEF cells for overlapping α-ESA/α-ESA-me candidate protein nodes and set of ferroptosis associated genes in RNA-Seq dataset. (B) Relative mRNA expression of key ferroptosis node genes in NS and S-Eto MEF cells following 24hr treatment with compounds. Error bars represent SEM for n = 4 for NS and n = 3 for S-Eto/treatment groups.

**Figure S10.**
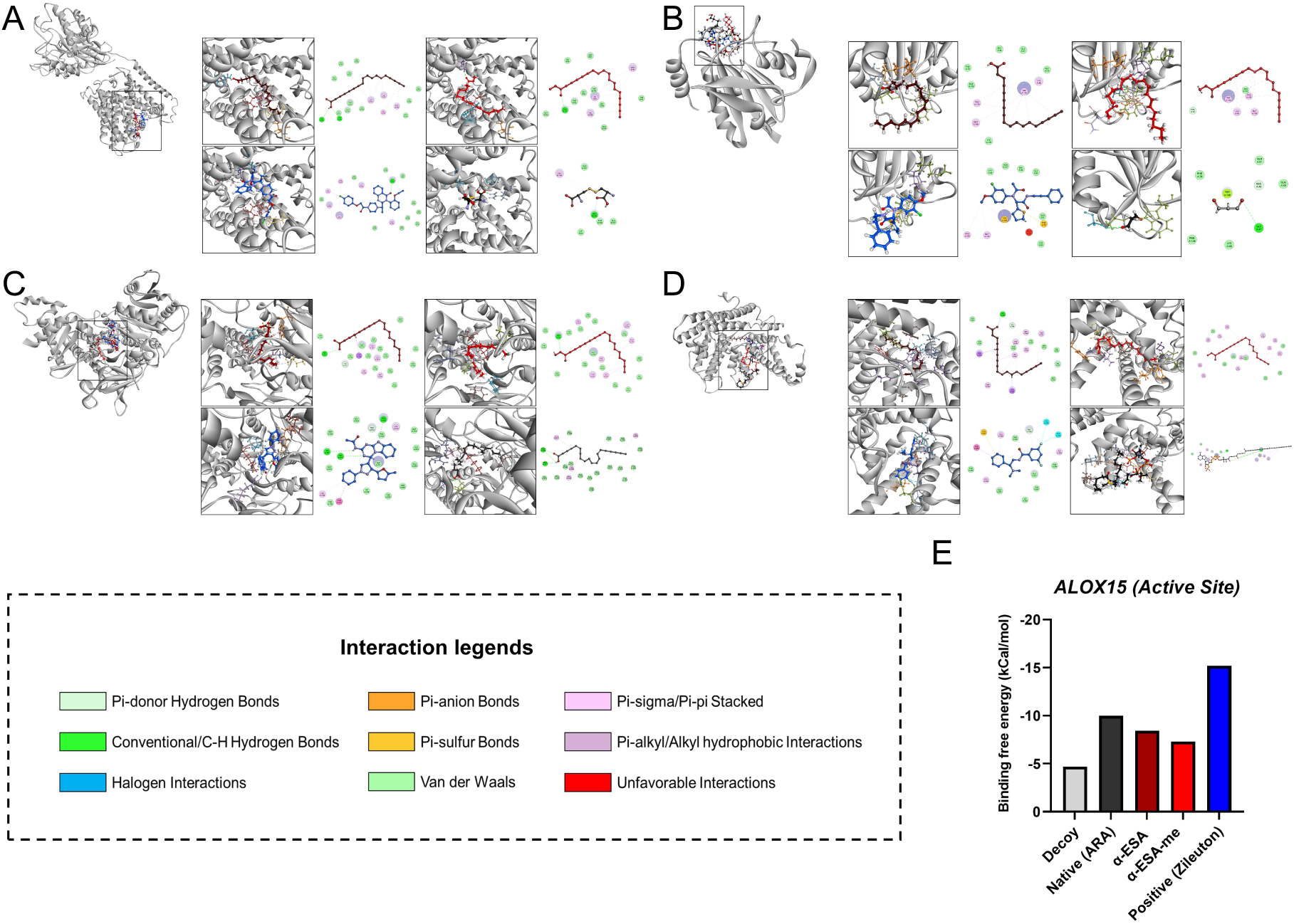
3D and 2D interaction plots of ligands in the substrate binding pocket/catalytic active site with neighboring pocket amino acid residues, related to Figure 7. Interaction plots for (A) System XC-, (B) GPX4, (C) ACSL4, and (D) LPCAT3. Ligand-residue interaction ligands are provided next to the plot. Compound categories follow color palette in (E). (E) Binding free energy scores of best-fitting conformers for each ligand within the ALOX15 substrate binding pocket. Lower binding free energies correspond to stronger ligand-receptor interactions.

